# PEP7 is a ligand for receptor kinase SIRK1 to regulate aquaporins and root growth

**DOI:** 10.1101/2022.04.05.486672

**Authors:** Jiahui Wang, Lin Xi, Xu Na Wu, Stefanie König, Leander Rohr, Theresia Neumann, Klaus Harter, Waltraud X. Schulze

**Affiliations:** Department of Plant Systems Biology, University of Hohenheim, 70593 Stuttgart, Germany; School of Life Science, Center for Life Sciences, Yunnan University, 650091 Kunming, People’s Republic of China; Center for Plant Molecular Biology, University of Tübingen, 72076 Tübingen, Germany

**Author notes:** Address of correspondence: Prof. Dr. Waltraud Schulze University of Hohenheim Department of Plant Systems Biology 70593 Stuttgart. equal contribution (Jiahui Wang, Lin Xi).

## Abstract

Plant receptor kinases constitute a large protein family that regulate various aspects of development and responses to external biotic and abiotic cues. Functional characterization of this protein family and particularly the identification of their ligands remains a major challenge in plant biology. Previously, we identified plasma membrane-intrinsic SUCROSE INDUCED RECEPTOR KINASE 1 (SIRK1) and QIAN SHOU KINASE 1 (QSK1) as a receptor / co-receptor pair involved in regulation of aquaporins in response to osmotic conditions induced by sucrose. Here, we identified a member of the Elicitor Peptide (PEP) family, namely PEP7, as the specific ligand of receptor kinase SIRK1. PEP7 binds to the extracellular domain of SIRK1 with a binding constant of 1.44±0.79 µM and is secreted to the apoplasm specifically in response to sucrose treatment. Stabilization of a signaling complex involving SIRK1, QSK1 and aquaporins as substrates is mediated by alterations in the external sucrose concentration or by PEP7 application. Moreover, the presence of PEP7 induces the phosphorylation of aquaporins *in vivo* and enhance water influx into protoplasts. The loss-of-function mutant of SIRK1 is not responsive to external PEP7 treatment regarding kinase activity, aquaporin phosphorylation and water influx activity. Our data indicate that the PEP7/SIRK1/QSK1 complex represents a crucial perception and response module mediating sucrose-controlled water flux in plants.

## Introduction

Plants as sessile organisms must be able to rapidly adapt to altering environmental conditions throughout the diurnal cycle and during their life span. This requires precise integration of extracellular information with intracellular (metabolic) signals. This integration of environmental and developmental signals in plants is often controlled by small peptides which then activate signaling cascades through receptor kinases. Receptor kinases constitute the biggest subclade within the plant kinome (Zulawski et al., 2014). The plant receptor kinases mainly serine/threonine kinases but show functional and structural relationships to receptor tyrosine kinases in animals (Shiu and Bleecker, 2001), and they are involved in plant developmental processes as well as in responses to biotic and abiotic cues (Osakabe et al., 2013).

The first peptide discovered in plants with signaling functions was systemin (Pearce et al., 1991), which at that time was postulated to also be perceived by a LRR-family receptor kinase (Scheer and Ryan, 2002). Since then, more and more signaling peptides were discovered, which are involved in cell to cell communication, developmental processes, and stress responses (Matsubayashi, 2014; Tavormina et al., 2015). It is a characteristic of the small signaling peptides that they are mobile between cells. Thus, the site of their secretion may be different from the site of their perception. For example, peptide CLAVATA3 is recognized by a LRR-receptor kinase expressed in neighboring cell files (Clark et al., 1997). The peptides can even be translocated throughout the plant, as exemplified by the CEP family peptides which are secreted by nitrogen starved roots, but their receptors are located in the shoot (Tabata et al., 2014).

These biologically active peptides are defined as being smaller than 100 amino acids and usually undergo a process of maturation from larger precursor proteins (Tavormina et al., 2015). Thus, the final active peptide is matured from its preproprotein through proteolytic processing by specific peptidases (Schaller et al., 2018). In many cases, this involves two steps, firstly the cleavage of an N-terminal signal sequence necessary for secretion of the protein to the apoplasm, and secondly release of the active peptide by cleavage of the prodomains. The peptides themselves are frequently subject to posttranslational modifications such as sulfatation, modification with sugar residues or hydroxyprolines (Matsubayashi, 2014). Formation of secondary structures through intramolecular disulfide bridges is a characteristic feature of cysteine-rich peptides, such as the RALF family (Matsubayashi, 2014). The process of peptide maturation has moved a variety of peptidase families into focus of attention in context of plant signaling pathways (Rautengarten et al., 2005). Subtilases were shown to be involved during maturation of the IDA peptide (Schardon et al., 2016) and also in phytosulfokine processing (Stührwohldt et al., 2021). Although it became apparent that receptor kinases are primary candidates for the recognition of the variety of biologically active peptides with signaling functions, for most of the receptor kinases the precise ligand remains unknown currently. In turn, also for many biologically active peptides, the receptors remain to be identified (Matsubayashi, 2003). Thus, for a more complete understanding of the functional implications of plant receptor kinases, it is of high interest to identify and characterize ligand–receptor pairs.

In the past, our group has studied sucrose-induced protein phosphorylation in a time course experiment resupplying sucrose to sucrose-starved *Arabidopsis* seedlings (Niittylä et al., 2007). Based on this time-course information, sucrose-dependent regulation of an aquaporin and sucrose-induced phosphorylation of the sucrose exporter SWEET11 by a protein complex involving SUCROSE-INDUCED RECEPTOR KINASE SIRK1 was discovered (Wu et al., 2013). As a follow-up to this work, we recently showed that the SIRK1 signaling complex is stabilized by the co-receptor QSK1 (Wu et al., 2019b). However, the ligand of this SIRK1-QSK1 receptor complex remained unknown. We therefore conducted a series of biochemical and physiological experiments to identify PEP7 as the specific ligand of receptor kinase SIRK1.

## Results

SUCROSE-INDUCED RECEPTOR KINASE (SIRK1) belongs to the LRR-receptor kinases (Zulawski et al., 2014). SIRK1 was found to be activated by external supply of sucrose (Wu et al., 2013). It interacts with co-receptor QIAN SHOU KINASE (QSK1) and in the active state regulates the opening status of aquaporins (Wu et al., 2019b). The majority of the LRR-receptor kinases for which a ligand is known so far, were found to bind small peptide ligands. Therefore, we followed the hypothesis that also SIRK1 could bind a small peptide ligand.

### Identification of SIRK1 ligand candidates

To systematically screen for a putative peptide ligand to SIRK1, we fractionated apoplasmic extracts derived from wild type liquid-grown *Arabidopsis* seedling cultures. The fractions were ultra-filtrated to exclude all protein components larger than 1 kDa. Each fraction was tested for its ability to *in vitro* induce the kinase activity of SIRK1-GFP, which was enriched from root tissue of hydroponic cultures two days after sucrose starvation (Figure 1A). Already kinase active SIRK1-GFP (SIRK1-GFPsuc) was enriched from sucrose stimulated hydroponic cultures (Wu et al., 2013) and used as a positive control (Figure 1B). Exposure of sucrose alone to SIRK1- GFP enriched from sucrose starved plants SIRK1-GFPstarv) was not able to induce SIRK1 activity (Figure 1B). In contrast, SIRK1-GFPstav activity was highly induced after exposure to protein fraction F2 and to a lesser extent to protein fraction F1. Kinase activity was lowest by exposure of SIRK1-GFPstarv to protein fraction F3 (Figure 1B). An aliquot of each fraction was analyzed by mass spectrometry. We performed two runs in parallel, one with standard protocol including trypsin digestion, one without prior tryptic digestion, in case the ligand candidates would not yield suitable tryptic peptides. Altogether, in fraction F1 we identified 1531 proteins, in fraction F2 4106 proteins, and in fraction F3 440 proteins. Among the identified proteins we then selected those candidate proteins which were present with higher spectral counts in fractions F1 and F2, and were not present in fraction F3. A total of 30 proteins were predicted to be secreted proteins, and 14 of these met the requirements of high abundance in fractions F1 and F2, but not in fraction F3 (Figure 1C). Two of these candidate proteins were also identified in the non-tryptic samples, namely RALF1 (AT1G02900) and PEP7 (AT5G09978). All tryptic and non-tryptic RALF1 and PEP7 peptides covered the C-terminal parts of the respective propeptides, which constitutes the biologically active RALF1 or PEP7 versions, respectively. Other RALF peptides such as RALF22, RALF23 and RALF33 were also identified in the fractions F1, F2 and F3, but did not strictly meet the criteria set for putative ligands as described above (Supplementary Table 1).

**Figure 1:**
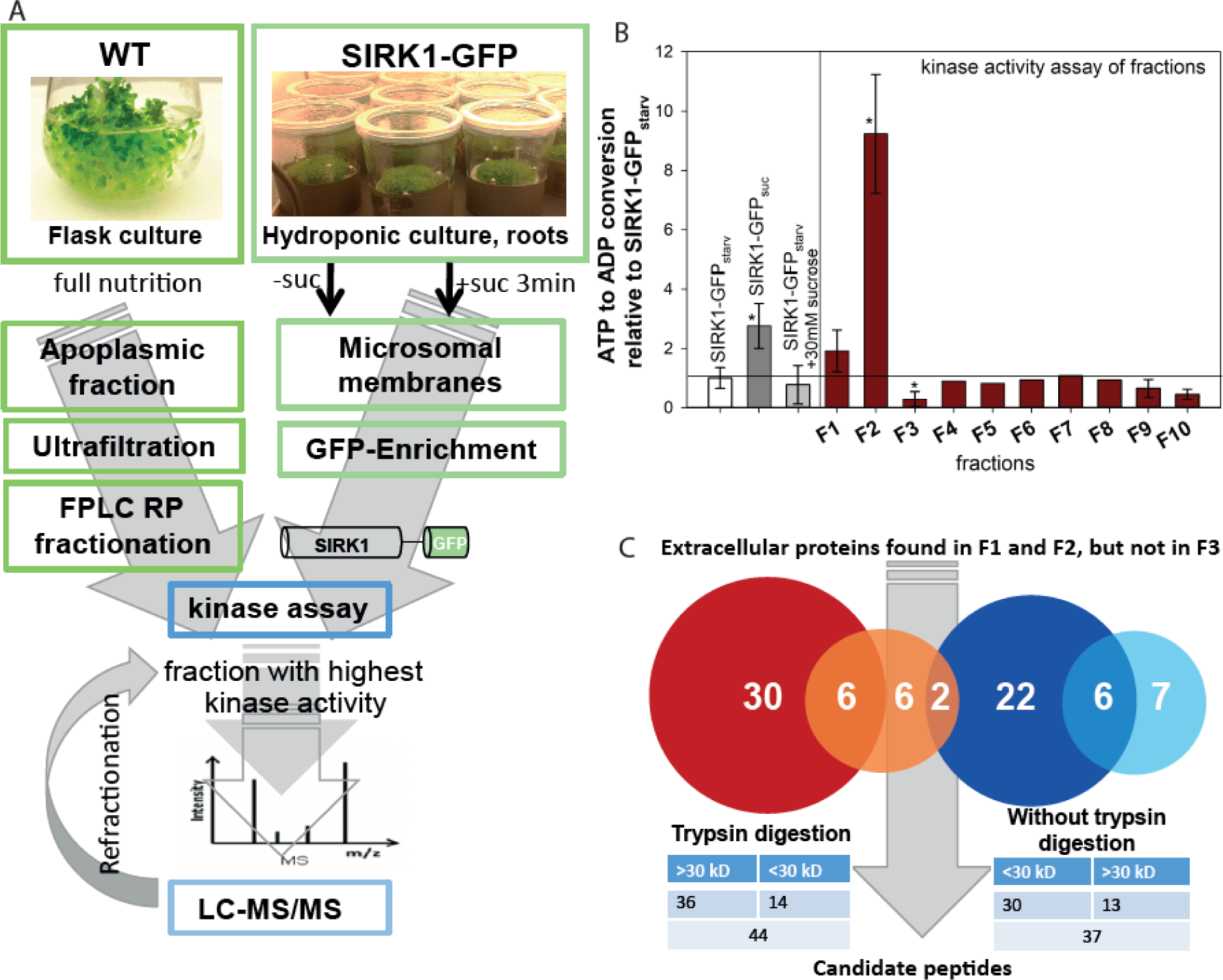
Screen for peptide ligands. (**A**) Workflow involving apoplasmic protein fractions exposed to SIRK1-GFP from hydroponic culture with and without sucrose stimulation. (**B**) Kinase activity of SIRK1- GFP exposed to different fractions of apoplasmic proteins. Asterisks indicate significant differences to SIRK1-GFP isolated from sucrose starved cultures (SIRK1-GFPstarv). For fractions F4 to F8 only one fraction was analyzed, all other fractions were tested in three independent replicates. (**C**) Summary of the number of proteins with predicted extracellular location identified in Fractions F1 and F2, but not F3.

We next used synthetic peptides of RALF1 and also RALF22, RALF32, and RALF33 at different concentrations to test for their ability to induce the kinase activity of SIRK1-GFPstarv (Supplementary Figure S1). Independent of the used concentrations, neither the RALF peptides, nor the non-related reference peptides IDA and CLE were able to induce SIRK1- GFPstarv kinase activity above background level (Supplementary Figure S1). In contrast, with the exception of RALF1, the application of the different peptides even reduced SIRK1-GFPstarv kinase activity compared to the control conditions (Supplementary Figure S1).

### PEP7 can bind to SIRK1 in biochemical and biophysical assays

The abundance of signaling peptide PEP7 increased in the apoplasm of liquid grown seedlings when resupplied with 30 mM sucrose after starvation (Figure 2A), but not when the sucrose-starved seedlings were supplied with related disaccharides such as sucralose or trehalose, or the monosaccharide glucose. Thus, the observed PEP7 accumulation was specific for sucrose-supplemented seedlings. To test, whether PEP7 has an effect on SIRK1-GFPstarv kinase activity, we performed *in vitro* dose-response assay. Increasing concentrations of synthetic PEP7 enhanced SIRK1-GFPstarv kinase activity, reaching saturation at PEP7 concentrations above 1 µM (Figure 2B). We then tested, whether PEP7 can directly bind to SIRK1-GFP. SIKR1-GFP – immobilized on anti-GFP magnetic beads (Supplementary Figure S2A) – was exposed to 1µM PEP7. After washing and elution, association of the putative ligand was detected by mass spectrometry. Indeed, SIKR1-GFP captured PEP7 from the solution (Figure 2C). Next, we performed a series of co-immunoprecipitation experiments to test for binding of PEP7 to the extracellular domain of SIRK1 (SIRK1-ECD). Firstly, StrepII-tagged SIRK1-ECD was purified after transient expression in *Nicotiana benthamiana* leaves (Supplementary Figure S2B) and immobilized to streptavidin beads and exposed to PEP7 at a concentration of 1µM. Again, PEP7 was detected by mass spectrometry among the eluted proteins which were bound to the bait (Figure 2D). In a reverse co-immunoprecipitation experiment, synthetic (His)6-tagged PEP7 was immobilized to Ni-NTA magnetic beads (Supplementary Figure S2C) and exposed either to 50 µg of SIRK1-GFP, 20 µg of SIRK1-ECD purified from *N. benthamiana* leaves after transient expression, or to root microsomal membrane preparations, which are expected to contain native full-length SIRK1 (Figure 2E). Indeed, in all three approaches, PEP7-(His)6 was found to capture SIRK1-GFP, StrepII-tagged SIRK1-ECD, or root-derived SIRK1 from the solutions. Next, we tested whether binding of SIRK1-ECD to immobilized PEP7-(His)6 can be competitively inhibited by addition of free, non-tagged PEP7. We detected a release of SIRK1- ECD from the PEP7-(His)6 beads with increasing concentrations of free PEP7 (Figure 2F). Saturation was reached at PEP7 concentrations above 1 µM. When free PEP6 (1 µM) was used to elute SIRK1-ECD from the PEP7-(His)6 beads, no release of SIRK1-ECD was observed. These experiments suggest that indeed PEP7 associates with SIRK1 *via* its ECD.

**Figure 2:**
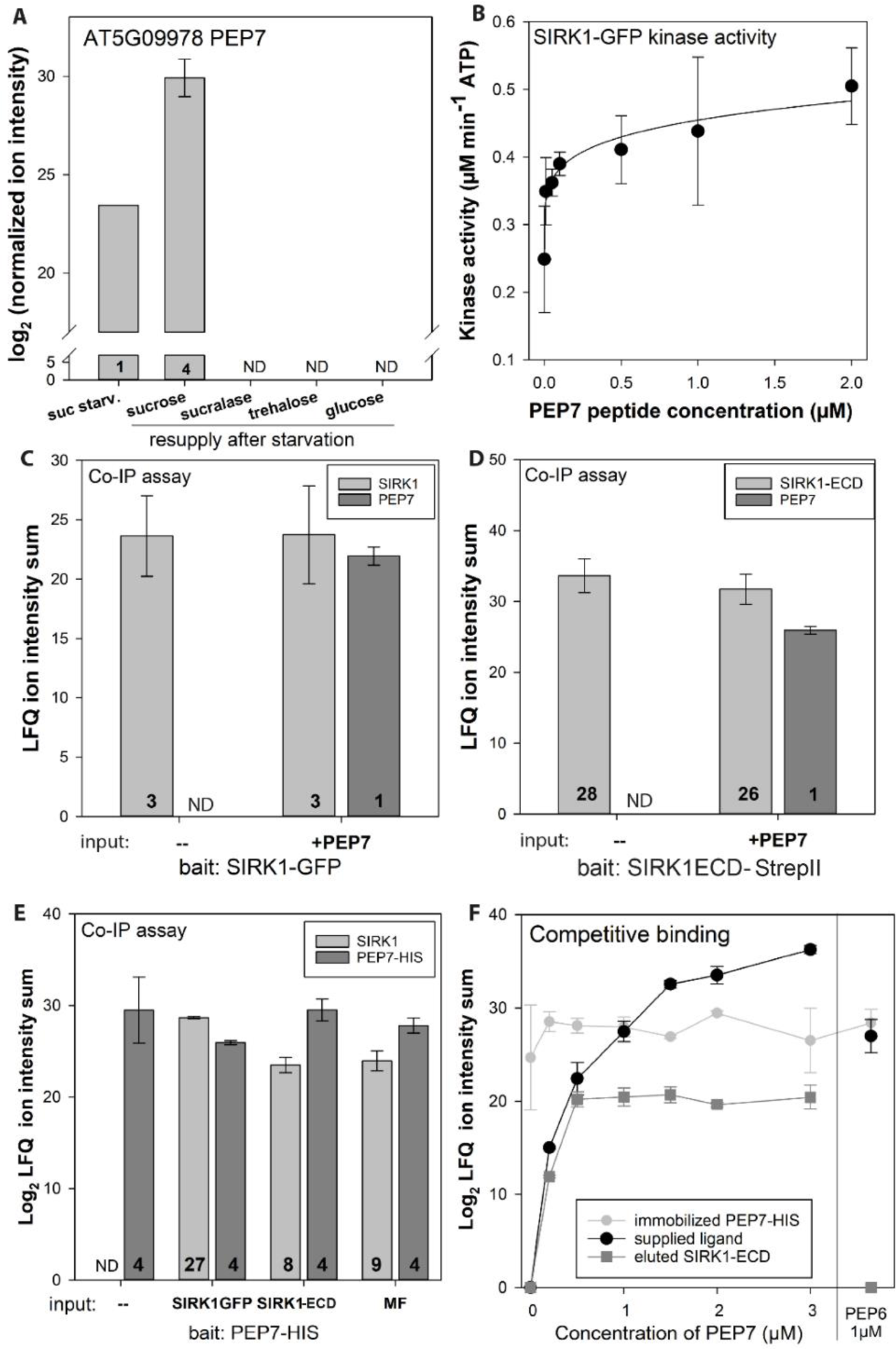
Characterization of PEP7 as a ligand candidate. (**A**) Normalized ion intensity of PEP7 in apoplasmic fractions in response to sucrose resupply or supply of other sugars. (**B**) Kinase activity of SIRK1-GFP induced by different concentrations of PEP7. (**C**) Normalized ion intensity (LFQ values) of SIRK1 and PEP7 in Co-IP assays with SIRK1-GFP with or without the supply of PEP7. (**D**) Normalized ion intensity (LFQ values) of SIRK1-ECD-HA and PEP7 in Co-IP assays with SIRK1-ECD-HA with or without the supply of PEP7. Averages of at least three biological replicates are displayed with standard deviations. In bar charts, numbers indicate the number of peptides identified. (**E**) Normalized LFQ- intensities of PEP7-HIS and SIRK1 in pulldowns with PEP7-HIS as a bait. (**F**) Competitive elution of SIRK1- ECD from pre-bound complexes with immobilized PEP7-HIS with different concentrations of PEP7. PEP6 was used as a control. In all panels, averages of at least three biological replicates are displayed with standard deviations. In bar charts, numbers indicate the number of peptides identified.

We used microscale thermophoresis to obtain more quantitative data with respect to the binding of PEP7 to SIRK1-ECD. StrepII-tagged SIRK1-ECD was purified (Supplementary Figure S3) and labeled with fluorescent dye RED-NHS. Incubation of different concentrations of PEP7 with SIRK1-ECD resulted in sigmoidal binding curve, allowing the calculation of the binding constant (Kd-value) of 1.44±0.79 µM PEP7 based on three assays from independent protein isolations (Figure 3). Adding approximate amount of co-receptor QSK1-ECD as a third component to the binding assay reduce the binding constant to a Kd of 57.01±22.23 nM. When PEP4 or PEP6 instead of PEP7 were used as putative ligands in the microscale thermophoresis assay, no sigmoidal dose-response curve could be obtained, and the signal-to-noise ratio (around 1.4) was too small to calculate a Kd-values. Thus, experimental evidence from the different association assays and microscale thermophoresis point to PEP7 being able to physically bind to SIRK1-ECD. QSK1-ECD could increase the binding affinity of SIRK1-ECD and PEP7, as expected from QSK1 function as a co-receptor. Thus, PEP7 is a strong candidate to be the specific ligand of SIRK1.

**Figure 3:**
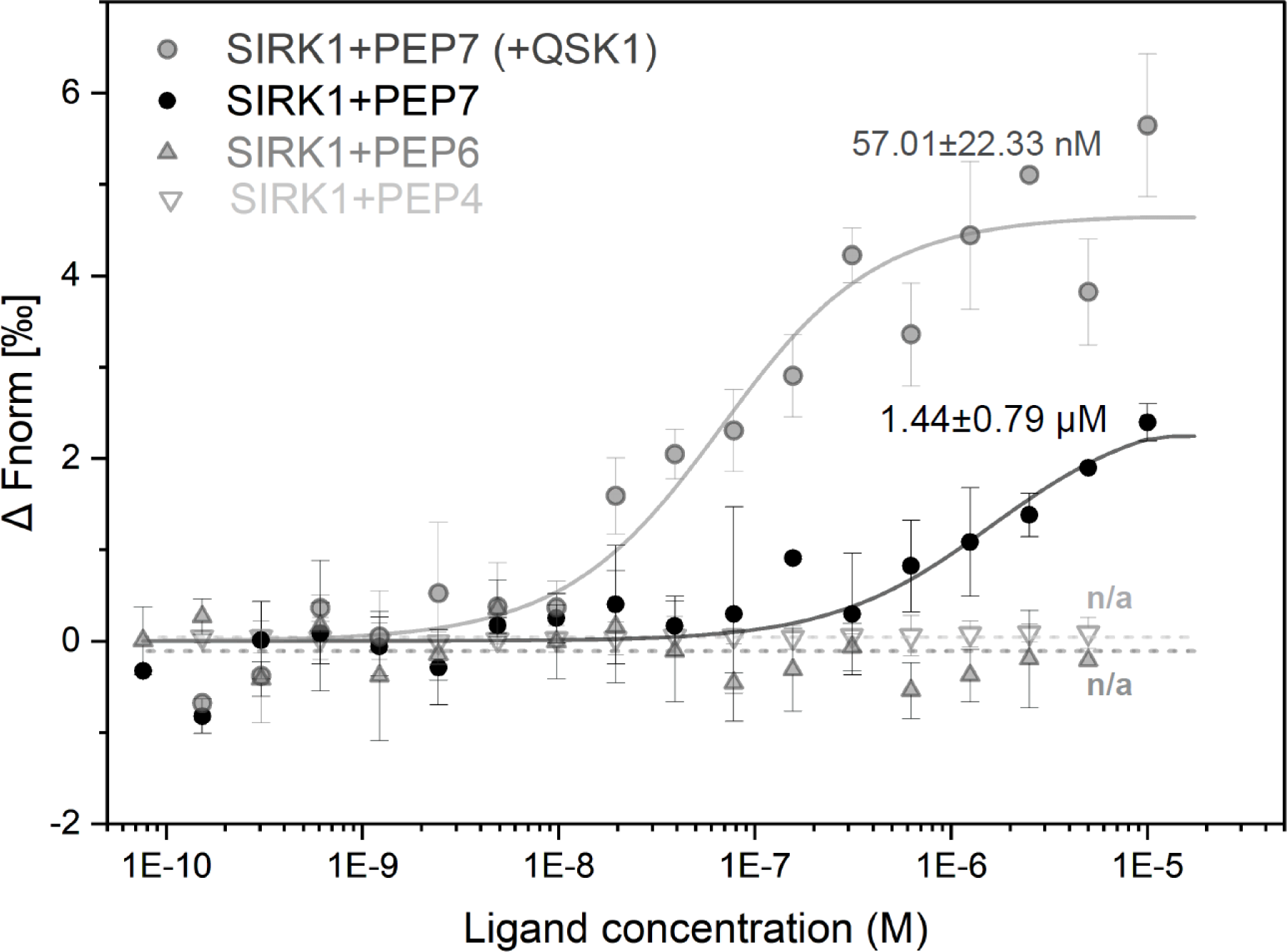
Microscale thermophoresis assay with the extracellular domain of SIRK1 (SIRK1-ECD) and PEP7 or PEP6 as putative ligands. In affinity measurement of SIRK1-ECD and PEP7 in presence of QSK1- ECD, 20nM QSK1-ECD was applied. A signal to noise ratio of 8 was achieved in assays for PEP7 as the ligand, while signal to noise ratio was 1.4 for PEP6 as the control. Kd values are as indicated. Each dot represents an average of at least three independent measurements. Error bars indicate SD. Fitting of the dose-response curve was performed by the instrument software.

### PEP7 induces the SIRK1 signaling complex and phosphorylation of SIRK1 substrates

SIRK1-GFP fusion expressed in the *sirk1* and *pep7* background was used to determine the interactome of SIRK1 after addition of sucrose, PEP7 or mock treatment following an established affinity purification protocols (Wu et al., 2019b) (Supplementary Table 2). Protein abundances were normalized to the protein abundance of the bait SIRK1-GFP (Figure 4A). Protein abundance ratios were calculated as log2 ratios of treatment (sucrose or PEP7) *versus* mock treatment (Figure 4B). Proteins, which showed a positive log2-fold change in their abundance, were considered to be recruited as interactors of SIRK1 in response to the respective treatment. Generally, there was a large overlap of induced log2-fold changes upon sucrose- or PEP7 treatment (Figure 4C) and we found a strong correlation of log2-fold changes in prey protein abundance induced by sucrose and PEP7 (Figure 4D). The known co-receptor QSK1 and its homolog QSK2 were found to be among the recruited interaction partners to SIRK1 in both treatments (Figure 4E). Also aquaporins were identified to be associated with SIRK1, again confirming earlier observations that SIRK1 interacts with aquaporins upon sucrose supply and are substrates for the SIRK1/QSK1 signaling complex (Wu et al., 2013; Wu et al., 2019b). (Figure 4F). Furthermore, PEP7 induced a comparable change of aquaporin abundance in the SIRK1-GFP pulldowns as sucrose (Figure 4F). We conclude that PEP7 induced the formation of a highly similar interactome of SIRK1 as was observed after treatment with external sucrose.

**Figure 4:**
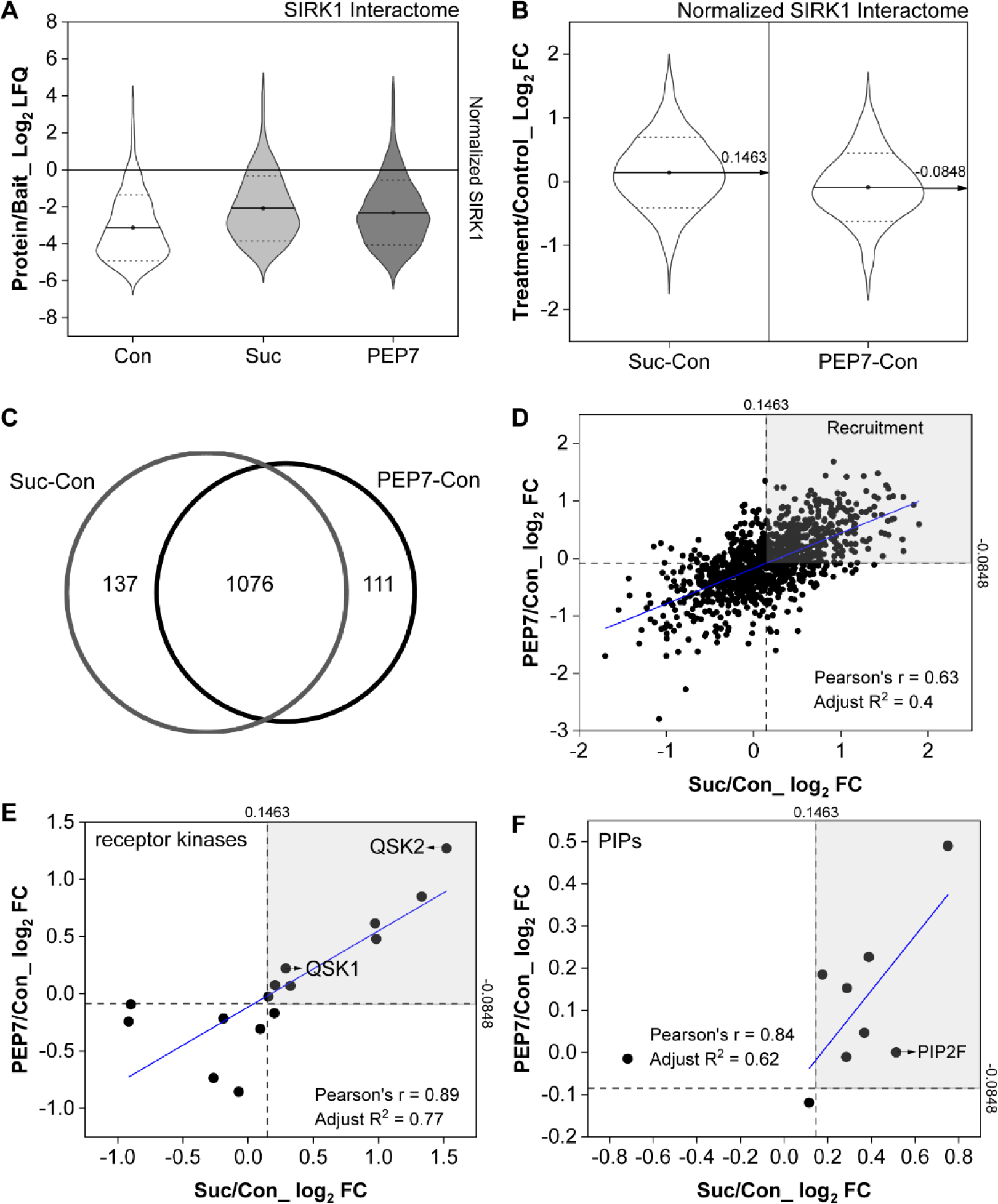
Interactome of SIRK1 induced by PEP7 and Sucrose (**A**) Protein abundance distribution normalized to bait (SIRK1) abundance. LFQ-values were taken as measure of protein abundance. (**B**) Distribution of treatment/control comparisons centered on median values. The two median values within each comparison are shown in the violin plot, respectively. (**C**) Venn diagram of two comparisons. (**D**) Correlation analysis of all protein comparisons under two treatments. Proteins with values greater than the median were considered to be recruited by SIRK1 under both treatments. (**E**) Correlation analysis of only receptor kinases and (**F**) Correlation analysis of only PIPs under two treatments.

As shown previously, the activation of SIRK1 by external sucrose supply (Wu et al., 2013; Wu et al., 2019b) involved the trans-phosphorylation of QSK1 and phosphorylation of the aquaporins (Figure 5A). In those experiments, hydroponic cultures of *Arabidopsis* plants where starved for sucrose and re-supplied with 30 mM sucrose solution for 5 minutes prior to harvesting of tissue and analysis of protein phosphorylation (Wu et al., 2013; Wu et al., 2019b). Here, we show that in wild type, also external supply of PEP7 (instead of sucrose) resulted in increased phosphorylation of QSK1/QSK2 and aquaporins (Figure 5B). The sucrose-induced phosphorylation of QSK1/QSK2 and aquaporins was reduced in the previously described *sirk1* loss-of-function mutant (Figure 5B). The external supply of PEP7 instead of sucrose to *sirk1* seedlings resulted in an even lower phosphorylation of QSK1/QSK2 and aquaporins compared to wild type. Moreover, the phosphorylation of aquaporins and QSK1/QSK2 was not observed in the *pep7* mutant upon sucrose supply. However, the increased phosphorylation of QSK1/QSK2 and aquaporins in *pep7* was restored by external supply of PEP7. In the *sirk1;pep7* double mutant, the phosphorylation of QSK1/QSK2 and aquaporins was neither induced by sucrose, nor rescued by external supply of PEP7. Importantly, the phosphopeptides (Supplementary Table 3) quantified for aquaporins corresponded to the known pore-gating phosphorylation sites from SoPIP2A (Figure 5C) suggesting that indeed, PEP7 induced pore opening phosphorylation of aquaporins similar as did treatment with external sucrose (Figure 5D).

**Figure 5:**
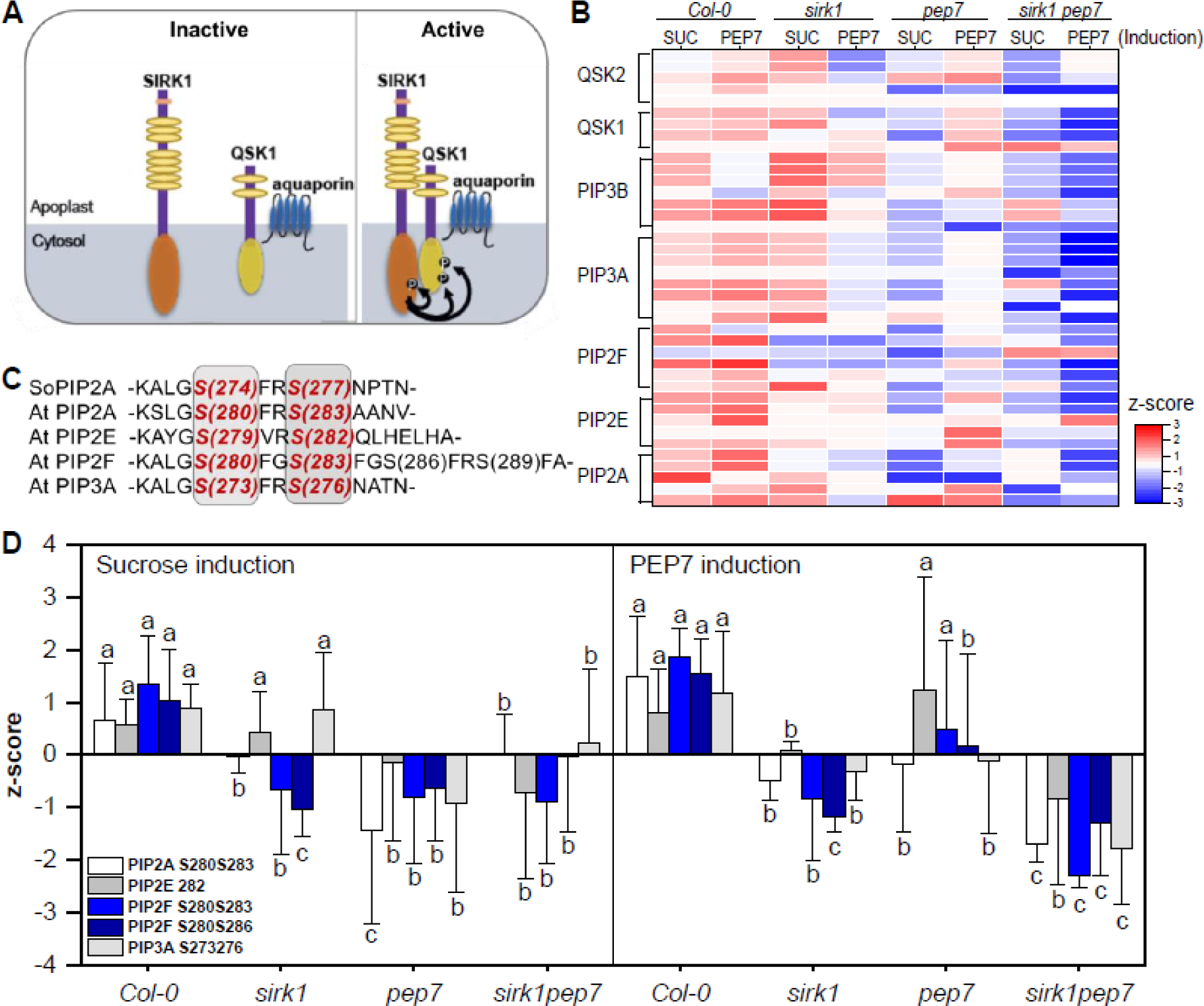
PEP7-induced alterations of phosphorylation of SIRK1 substrates. (**A**) SIRK1 signaling scheme highlighting substrates that presented previously. (**B**) Heatmap of means of z-scored phosphosite intensities of different phosphopeptides identified for known members of the SIRK1 signaling pathway. Each row represents one phosphopeptide. (**C**) Sequence alignment of C-terminal phosphorylated plant aquaporins. Grey area indicates the conserved residues that can be phosphorylated. (**D**) z-scored phosphorylation levels of conserved residues of PIPs that correspond to the pore gating that induced by sucrose and PEP7. Small letters indicate significant differences (p<0.05; pairwise t-test) for each phosphopeptide between genotypes.

### PEP7 may in vivo stabilize association of SIRK1 and QSK1 in preformed complexes

To investigate potential effect of PEP7 on SIRK1 and QSK1 associations, we measured their spatial proximity *in vivo* by Förster resonance energy transfer by fluorescence lifetime (FLT) imaging (FRET-FLIM). QSK1-GFP and SIRK1-mCherry co-localize in the plasma membrane with or without the supply of PEP7 in transiently transformed *N. benthamiana* epidermal leaf cells (Supplementary Figure S4A, B). To determine the background FRET-FLIM values, the FRET donor QSK1-GFP was expressed alone and the fluorescence life time (FLT) determined. In all cases, where QSK1-GFP and SIRK1-mCherry (FRET acceptor) were co-expressed, a reduction in FLT was observed indicating a close spatial proximity (less than 10 nm (Glöckner et al., 2020)) of the two proteins. There was no significant further reduction in FLT when PEP7 was applied compared to mock treatment (Supplementary Figure S4B). To exclude the possibility that endogenous PEP7 may mask the effects of externally applied PEP7 on QSK1-GFP/SIRK1- mCherry association, we performed the FRET-FLIM experiments in the presence of the protease inhibitor leupeptin. Leupeptin inhibits the metacaspase 4 (MC4) (Vercammen et al., 2004), which releases PEP7 from its propetide (Shen et al., 2019). After treatment with leupeptin, the FLT of QSK1-GFP in the donor-only control was reduced in response to PEP7 addition (Supplementary Figure S4B). The FLT of QSK1-GFP in the presence of SIRK1-mCherry was not significantly altered in response to PEP7 application even after treatment with leupeptin (Supplementary Figure S4B). Likely, SIRK1 and QSK1 are associated in preformed nano-structured membrane domains in the absence of PEP7 as it is described for several other LRR-RK complexes (Bucherl et al., 2017; Glöckner et al., 2020; Gronnier et al., 2022). Therefore, PEP7 may act as a stabilizing factor of the QSK1/SIRK1 association, thereby transforming the pre-formed inactive complexes into active ones.

### PEP7 affects water influx to protoplasts via receptor kinase SIRK1

Sucrose as an osmotic agent was previously shown to induce water influx into protoplasts, and this sucrose-induced water influx was significantly reduced in the *sirk1* mutant (Wu et al., 2013). We performed protoplast swelling assays with wild type, *sirk1* and *pep7* single mutants as well as the *sirk1;pep7* double mutant to test if supply of PEP7 was also able to induce water influx and whether this was dependent on the presence of SIRK1 receptor kinase. In wild type, mannitol and sucrose induced water influx across the plasma membrane (Figure 6A, B). External supply of PEP7 in wild type significantly increased water influx density compared to mannitol or as observed in presence of sucrose. In the *sirk1* mutant, mannitol induced water influx, but sucrose supply did not. Supply of PEP7 to *sirk1* mutant did not increase water influx density (Figure 6A, B), which remained low, similar to water flux density observed in *sirk1* with sucrose treatment. In the *pep7* mutant, mannitol induced water influx, but sucrose supply resulted in low water influx density, similar as in the *sirk1* mutant. Strikingly, external supply of PEP7 to the *pep7* mutant restored water influx to water flux densities not different from wild type (Figure 6A, B). In the *sirk1;pep7* double mutant, neither sucrose nor PEP7 supply resulted in enhanced water influx densities, suggesting that SIRK1 is required for the swelling response.

**Figure 6:**
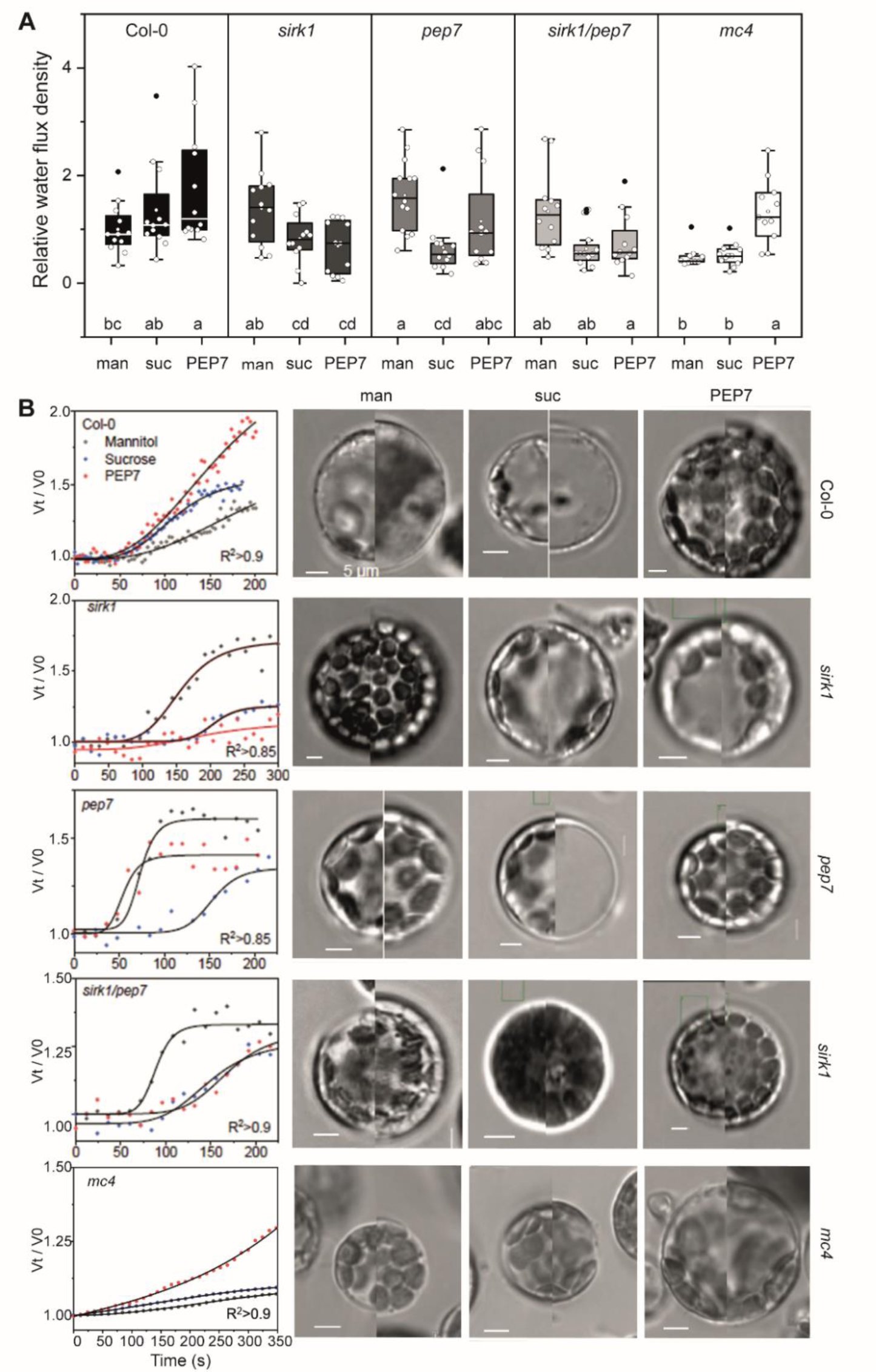
Water influx density of protoplasts induced by osmotic changes through mannitol, sucrose, or by the supply of PEP7 in wild type (Col-0), *sirk1*, *pep7*, the double mutant *sirk1;pep7* and *mc4*. (**A**) Boxplots show the water flux density relative to wild type. The average of Col-0 under mannitol treatment was used as the control and set to 1. Vertical lines in the boxes indicate the median and upper/lower borders represent the 25^th^ percentile. White dots represent individual measurements. Small letters indicate significant differences (p<0.05) between treatments as determined by oneway ANOVA with Holm-Sidak correction. (B) Volume change of protoplasts over time under induced by mannitol only, in presence of sucrose or in presence of PEP7 and representative images of protoplasts at high and low osmolarity conditions. Scale bar 5µm. man: mannitol; suc: sucrose.

Recently, the MC4 was identified to be involved in PEP7 maturation (Shen et al., 2019). Interestingly, water influx rates of the *mc4* loss-of function mutant resembled the water influx rates of the *pep7* mutant. In *mc4*, as in *pep7*, sucrose did not induce water influx, but PEP7 also in *mc4* was able to restore water influx to the protoplasts (Figure 6A, B). This further supports our conclusion of PEP7, as being produced by MC4 activity, to be involved in regulation of water influx.

We then tested whether other members of the PEP-family were able to affect water influx to protoplasts in a similar way. PEP4 and PEP6 were used since they were also found to be expressed in root tissue (Shulse et al., 2019). Water influx densities in wild type were not affected by the presence of PEP4 or PEP6. When the receptor kinase SIRK1 was absent (*sirk1* mutant), water influx densities in presence of PEP4 or PEP6 were not different from wild type suggesting that PEP4 and PEP6 did not require the presence of receptor kinase SIRK1 (Supplementary Figure S5). Moreover, in the *pep7* mutant, no difference in water influx densities compared to wild type was observed when PEP4 or PEP6 were supplied externally (Supplementary Figure S5).

### Root growth phenotype induced by PEP7 require receptor kinase SIRK1

Next, we explored the role of the PEP7/SIRK1 signaling pathway on organismal level. We hypothesize that root growth, which relies on sucrose supply as carbon source and water transport for turgor buildup (Péret et al., 2012), may be impaired in the *sirk1* mutant. Indeed, after six days of growth at low external sucrose supply (0.02% or 0.2%), the primary root of *sirk1* was significantly shorter than in wild type. This short root phenotype was enhanced in the *sirk1 qsk1* double mutant (Figure 7A, D), which is expected when both, receptor (SIRK1) and co-receptor (QSK1) (Wu et al., 2019b) are non-functional. On agar plates with high (1% or 2%) external sucrose supply, *sirk1* as well as *sirk1 qsk1* exhibited no significant difference in root length compared to wild type (Figure 7B, D, Supplementary Figure S6). Likely the high external sucrose provides high enough osmotic potential to induce cell expansion independently of SIRK1. The short root phenotype of *sirk1* under low sucrose conditions could be rescued by over-expressing SIRK1 (Supplementary Figure S6). Thus, the root growth phenotype of *sirk1* was dependent on SIRK1 expression and activity. Interestingly, the *pep7* mutant also showed a significantly shorter root growth under low sucrose (Figure 7A, D) and root growth under external sucrose supply was slightly larger than in wild type (Figure 7B, D).

**Figure 7:**
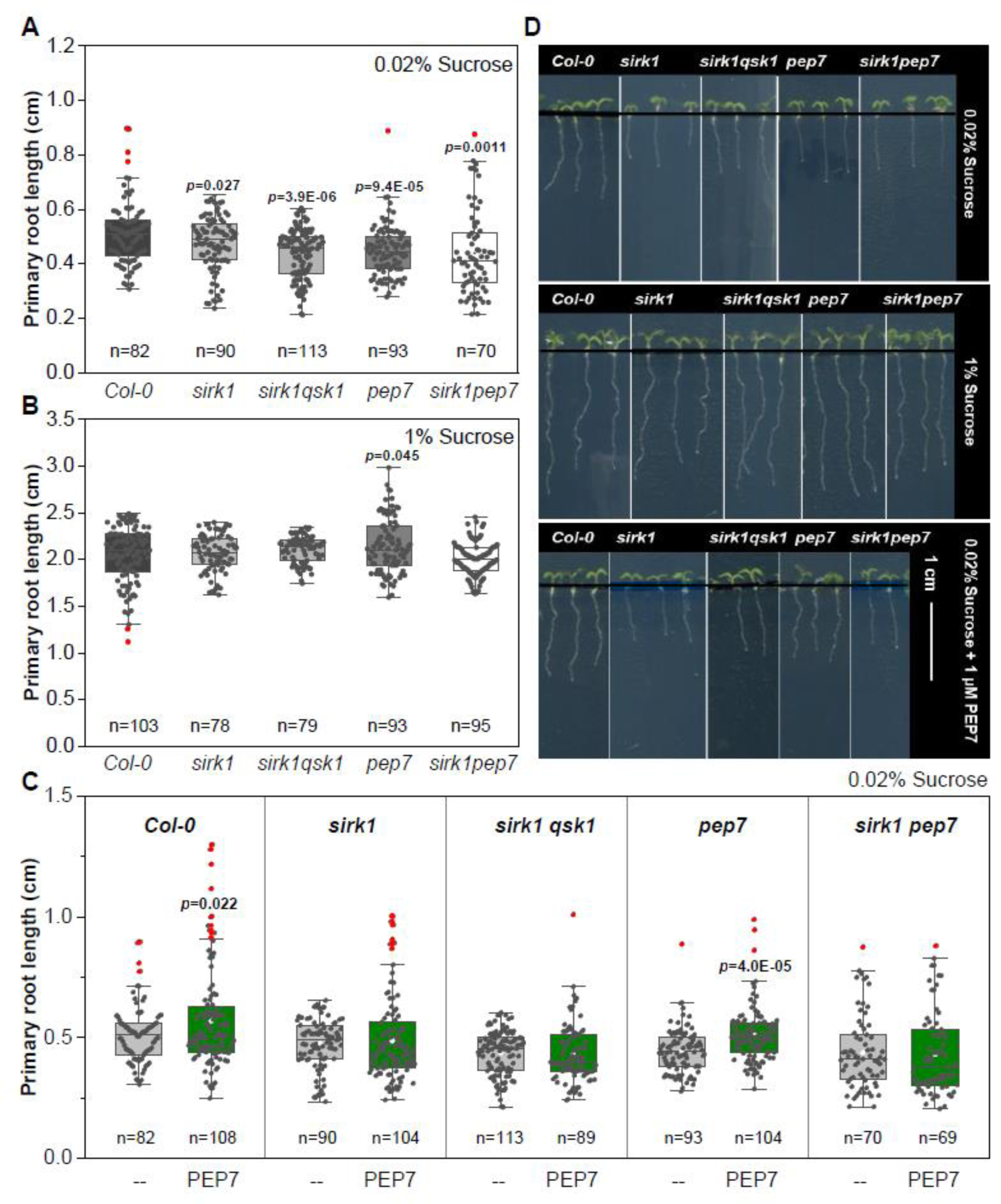
Root growth phenotype. Wild type (Col-0), *sirk1*, *pep7*, and *sirk1 qsk1* and *sirk1 pep7* double mutants grown on agar plates of (**A**) low external sucrose and (**B**) high external sucrose. (**C**) The effect of PEP7 supply (1µM) to roots was tested for the four genotypes at low external sucrose. (**D**) Scanned images of roots. Boxplots represent values of individual plants, numbers of plants analyzed are noted below each box. In panels A and B, p-values indicate significant differences to wild type. In panel C, p- values indicate significant differences of the PEP7-treatment to control treatment within each genotype.

Supply of external PEP7 (1 µM) to agar plates with low sucrose content slightly increased the root growth in wild type compared to control conditions (Figure 7C, D). In the *sirk1* mutant, external supply of PEP7 did not affect the short-root phenotype. Also in the receptor/co-receptor mutant *sirk1;qsk1*, external supply of PEP7 had no effect. In the *pep7* mutant, supply of external PEP7 restored the primary root length to wild type levels (Figure 7C, D), but this was not observed in the *sirk1;pep7* mutant in which also the receptor SIRK1 was missing. The root-growth enhancing effect of external PEP7 supply was not observed in *sirk1*; *sirk1;qsk1* and *sirk1;pep7* mutants. Again, these results suggest that PEP7 signaling requires presence of the SIRK1 receptor.

## Discussion

PEP7 belongs to a family of “danger signaling peptides” consisting of 8 family members (Zhang et al., 2016a). Out of this family, PEP1 was shown to be recognized by receptors PEPR1 (AT1G73080) and PEPR2 (AT1G17750) (Krol et al., 2010; Yamaguchi et al., 2010), but very little is known about the other PEP-family members. Here, we point to a signaling pathway for PEP7. Based on the sequences of the active peptides, PEP6 shows strongest similarity to PEP7 and therefore was used as a control. PEP4, although its propeptide shares expression with PEP7 in root tissue, shows strongest dissimiliarity to PEP7 and was used as a second negative control.

### PEP7 as a member of the danger-signaling peptide family

PROPEP7 is expressed in the root elongation zone and lateral root primordia (Bartels et al., 2013). The expression does not show overlaps with expression of the known PEP-receptors PEPR1 and PEPR2 (Bartels et al., 2013) suggesting a different perception pathway for PEP7. SIRK1 shows unique expression patterns compared to PEPR1 and PEPR2 especially in sink tissues of shoot and root (Winter et al., 2007). However, the SIRK1 signaling pathway is linked with components of PEPR1 and PEPR2 signaling through common interaction partners. For instance, BAK1, two HERK2 paralogs and QSK1 – all classified as co-receptors (Xi et al., 2019) - are suggested as common interaction partners for all three receptors (PEPR1, PEPR2, SIRK1) (Arabidopsis Interactome Mapping Consortium, 2011; Jones et al., 2014).

PROPEP7 was found by single-cell sequencing of root cells, clustering with marker genes of mature root hairs (Ryu et al., 2019). In the same study, PROPEP6 was found in a gene cluster with marker genes of the endocortex, while other members of the PEP-family or the respective PEPR receptors were not found. Most interestingly, in another single-cell sequencing study of root cells (Shulse et al., 2019), expression of PROPEP7 was induced by external sucrose supply, while expression of PROPEP6 and PROPEP4 were not induced by sucrose (Shulse et al., 2019). Thus, members of the PEP-family seem to be involved also in other signaling functions besides biotic stress responses, and were shown to be released also upon changes in plant central carbon metabolism. Our research points to an involvement of PEP7 in regulatory processes induces by osmotic imbalances in sucrose availability involving SIRK1 as a receptor.

### Processing of PEP7

Recently, type-II metacaspases were found to be involved in processing of the plant elicitor peptides (Shen et al., 2019), specifically Metacaspase 4 (AT1G79340) was able to process also PROPEP7 by releasing the active peptide from the C-terminus of the propeptide after cleavage after arginine or lysine. The processing of PEPs was induced by calcium signals (Hander et al., 2019), and required intramolecular proteolysis even of the metacaspase. The mRNA of MC4 is cell-to-cell mobile (Thieme et al., 2015). Little is known about further requirements for processing of the PEP precursors. Our work supports that MC4 may indeed be involved in PEP7 maturation as concluded from the protoplast swelling assays of the *mc4* mutant with supply of external PEP7 and/or sucrose (Figure 6).

### Receptor-ligand binding

Receptor kinases are known to have strong binding affinities for their ligands. Brassinolide binds to the BRI1 receptor with a Kd of 15 nM as determined by immunoprecipitation assays (Wang et al., 2001). Scatchard plots suggested the Kd of CLAVATA3 binding to its receptor CLAVATA1 at 17 nM (Ogawa et al., 2008). The binding of CLE41/44 to receptor PXY with Kd of 33 nM was detected by using isothermal titration calorimetry (Zhang et al., 2016b). The binding of RALF to its receptor FERONIA with a Kd of around 1 µM was determined by microscale thermophoresis. Interestingly, far higher binding constants were found for binding of the peptide IDA to its receptor HASEA with a Kd of 20 µM using isothermal titration calorimetry (Santiago et al., 2016). The higher Kd values, as in the case of IDA, were obtained under conditions in which the co-receptor (SERK1) was not present. In that regards, the Kd-values obtained here for the binding of PEP7 to receptor SIRK1 are within this range at around 1µM. Strikingly, by adding the co-receptor QSK1 to the binding assay a significant higher binding affinity for PEP7 was observed (Kd = 57.01±22.3).

One important function of co-receptors is in the stabilization of the interaction of the receptor with the ligand. The co-receptor contributes as a shape-complementary component and by interaction with the receptor holds the ligand in place (Santiago et al., 2013; Sun et al., 2013a; Sun et al., 2013b; Zhang et al., 2016b; Zhang et al., 2016c). We use the approximated same amount of QSK1-ECD as the SIRK1-ECD in this assay, but it is not yet known if different ratios of receptor and co-receptor could result in different Kd of SIRK1and PEP7. Thus, our biochemical assays, the proteomics study of protein complexes and their phosphorylation, as well as the cellular and whole plant responses support PEP7 as a ligand to receptor kinase SIRK1.

### Preformed complexes of SIRK1 and QSK1

Our results suggest that PEP7 functions as the ligand for the nano-structured SIRK1/QSK1 complex and induces SIRK1-dependent signaling. Apparently, PEP7 is produced in conditions of changed sucrose supply. According to our data, the ligand PEP7 would induce the formation of the SIRK1 signaling complex. However, our attempts to visualize the recruitment of co-receptor QSK1 to ligand-activated SIRK1 using FLIM/FRET experiments revealed no clear differences between treatments with PEP7 or controls. This supports the hypothesis that receptor and co-receptor are arranged in pre-formed complexes within nano-domains (Bucherl et al., 2017; Gronnier et al., 2022) and that these complexes are likely stabilized or re-arranged by the ligand to reveal their full activity (Caesar et al., 2011; Ladwig et al., 2015).

Our finding that SIRK1 and QSK1 are found with higher abundance in pull-down experiments after treatment with PEP7 support the view of complex stabilization so that it can be detected after purification. The phosphorylation of scaffold proteins may be important in formation and re-arrangement of such preformed signaling domains (Perraki et al., 2018). In any case, binding of PEP7 to the SIRK1-QSK1 protein complex leads to activation of downstream signaling cascade involving activation of aquaporins and subsequent changes in water influx to the cell.

### PEP7 and the response to sucrose

Our findings strongly point to a receptor/co-receptor signaling pathway of SIRK1/QSK1 which is activated by the Elicitor Peptide PEP7 and results in opening of aquaporins by SIRK1- dependent phosphorylation. The opening status of aquaporins was shown to also have effects on lateral root emergence (Péret et al., 2012) *via* auxin signaling, and root hydraulic conductivity is affected by aquaporin abundance, even in response to signals from the shoot (Vandeleur et al., 2014). Thus, effects of the SIRK1/QSK1/PEP7 signaling pathway on root growth may also be indirect through reduced cell expansion in mutants where aquaporin activation is impaired.

## Conclusion

We here show that PEP7 is secreted in response to external sucrose supply. Thus, activation of SIRK1 by sucrose, as studied in previous work (Wu et al., 2019a; Wu et al., 2013), requires sensing of sucrose status and maturation of PEP7 before the receptor SIRK1 is activated. A challenging question remains for further study, as to whether external sucrose triggers the PEP7 maturation, or whether sucrose acts after uptake into the cells.

## Materials and Methods

### Plant Materials

*Arabidopsis* seeds of wild type (col-0), *sirk1* (SALK_125543), *qsk1* (SALK_019840), *pep7* (SALK_025824), *pep6* (SALK_141703) and *mc4* (SAIL_856_D05) were used. Double mutants *sirk1;pep7* and *sirk1;qsk1* were obtained by crossings of the respective single mutants. Furthermore, we overexpressed of SIRK1 (*35S::SIRK1-GFP*) in the background of *sirk1* and *pep7*. Homozygous T-DNA insertional mutants *sirk1* and double mutant *sirk1;qsk1* were confirmed *via* PCR amplification as reported previously (Wu et al., 2019b). Mutants of *pep7* and *sirk1;pep7* were confirmed *via* PCR amplification using T-DNA border primer LBb1.3 (5’-ATTTTGCCGATTTCGGAAC-3’) and gene-specific primers (PEP7-RP: 5’- GGAAGGTGCCTAGTTGGTACC -3’, PEP7-LP: 5’-GTTTTCACGTTTCAAATTCGG-3’; SIRK1-RP: 5’- TTTCCAGCATTTCCAACACTC-3’, SIRK1-LP: 5’-CACTAAGCTTGTTGAGGTCGC-3’) (Supplementary Figure S7).

### Hydroponic cultures

Plants were germinated and grown under 16/8 day/night (22 °C, 120 µE/s·m^2^) in ½MS medium plus 0.5% sucrose in a hydroponic cultivation system (Schlesier et al., 2003). Whole seedling cultures were grown in 50 mL of ½MS medium with 0.5% sucrose as described (Niittylä et al., 2007). Sucrose starvation-resupply experiments were performed as described (Niittylä et al., 2007). Plant material was collected after sucrose starvation (no resupply), after sucrose treatment (1% sucrose for 5 min), or PEP7 treatment (1 µM PEP7 for 5 min). Addition of glucose (33 mM), sucralose (30 mM) and trehalose (40 mM) with the same osmolarity (Vapro 5600, Wescor Biomedical Systems) as the 30 mM sucrose solution were used as control treatments in some experiments.

### Protein extraction and ultrafiltration

At harvesting, tissue was flash-frozen in liquid nitrogen. After breaking 300 g of frozen tissue to coarse pieces, the frozen tissue was transferred to a glass grinder (GLASS/PTFE Potter Elvehjem Tissue Grinder 30 ml) and 0.1% trifluoroacetic acid (TFA) was added to a volume of about 30 mL per sample. After tissue homogenization, the solution was filtered through four layers of gaze (Miracloth, Mecrk Millipore). Remaining cell debris was pelleted at 4°C and 10000 g for 15 minutes. The supernatant was used for further ultrafiltration and fractionation. The protein extract was firstly filtered through a 0.45 µm filter to get rid of any insolubilized material. Afterwards, the protein extract was concentrated over a tangential flow filtration (Minimate^TM^ Tangantial Flow Filtration Systems, Pall Corporation) using a molecular weight cutoff of 1 kDa (i.e. retaining anything larger than 1 kDa).

### Fractionation by size-exclusion chromatography

The ultrafiltrate was fractioned by FPLC (NGC chromatography systems, BioRad) by reverse phase chromatography using a Bio-scale MT column (2 mL, BioRad). The column was packed with Macro-Prep® t-Butyl and Methyl Hydrophobic Interaction Chromatography Media. Before fractionation, the FPLC system was washed with 20 % ethanol with a flowrate of 10 ml·min^-1^ (50% pump A/50 % pump B). The system was then washed with 2.5 ml H2O at a speed of 0.5 ml min^-1^ for 5 min. The column was equilibrated with 6 ml 2% acetonitrile and 0.1 % TFA (8.57 min x 0.7 ml·min^-1^) and 500 µL sample was loaded to the column. A linear elution gradient was run from 100% pump A to 100% pump B. Pump A was supplied with 2% acetonitrile with 0.1 % TFA and pump B with 50 % acetonitrile with 0.1 % TFA. The gradient was run for 28 min of which the last 3 min were 100% pump B. The fractioned solution was collected with a fraction collector (BioFrac ^TM^ Fraction Collector, BioRad) in ten 2 ml fractions.

### Transient expression of recombinant SIRK1 in N. benthamiana

*Agrobacterium tumefaciens* GV3101 strain harboring the relevant constructs and strain C58C1 (pBinG1 vector with P19 gene) were grown in liquid LB medium (10 gl^-1^ Tryptone, 5 gl^-1^ yeast extract and 10 gl^-1^ NaCl) at 28°C with appropriate antibiotics. For infiltration, *Agrobacterium* culture were adjusted to final OD600 0.5 for GV3101 and 0.25 for strain C58C1 in a mixture, using infiltration buffer 10 mM MES-KOH pH 5.6, 200 µM Acetosyringon, 10 mM MgCl2. 3-4 week-old *N. benthamiana* leaves were syringe infiltrated and harvested after 48 h. The harvested leaves were flash-frozen in liquid nitrogen before protein extraction and purification. SIRK1-ECD and QSK1-ECD were expressed as C-terminal fusion with an HA and StrepII tag using the pXCS vector series, which expressed in GV3101::pMP90RK strain (Witte et al., 2004).

### Protein extraction and Strep-Tag purification

*N. benthamiana* leaf material harboring SIRK1- ECD or QSK1-ECD was ground in liquid nitrogen (Witte et al., 2004) and thawed in 100 mM Tris-HCl pH 8.0, 150 mM NaCl, 1 mM EDTA, 2% w/v PVPP, 0.5% Triton X-100, 1 mM PMSF and 5 mM DTT. The mixture was incubated at 4°C for 2 h under constant shaking and after centrifugation (14000 rpm, 10 min, 4 °C) the supernatant was subjected into equilibrated Strep-Tactin® column (2-4013-001, IBA GmbH) according to the manufacturer tutorial. The StrepII tagged protein were eluted with 50 mM biotin, 100 mM Tris-HCl pH 8.0, 150 mM NaCl, 1 mM EDTA. Eluates were passed through a Macrosep Advance Centrifugal Devices with Omega Membrane 10K (MAP010C36, Pall Corporation) to get rid of the biotin contamination in the eluate. The target protein (SIRK1-ECD and QSK1-ECD) was collected from the sample reservoir.

### Microsomal membrane preparation

Microsomal membranes were enriched by differential centrifugation as described (Pertl et al., 2001; Wu et al., 2019a; Wu et al., 2013). 1.5 g of fresh tissue was homogenized in 10 ml ice-cold 330mM mannitol, 100 mM KCl, 1 mM EDTA, 50 mM Tris-MES pH 7.5, 5 mM DTT, 1 mM PMSF and 0.5% v/v Protease inhibitor cocktail (P9599, Sigma-Aldrich) and phosphatase inhibitors (25 mM NaF, 1 mM Na3VO4, 1 mM benzamidin, 3 µM leupeptin). The homogenate was centrifuged for 15 minutes at 7500 × g at 4 °C. The supernatant was centrifuged again for 75 minutes at 48,000 × g at 4 °C resulting in the microsomal membrane pellet.

### Pull-downs of GFP-tagged SIRK1

Microsomal proteins (100 µg) resuspended in 330 mM mannitol, 25 mM Tris-MES pH 7.5, 0.5 mM DTT was incubated with 25 µl of anti-GFP agarose beads (Chromotek) (Wu et al., 2013). After incubation, beads were collected and washed twice with 500 µl 10 mM Tris-HCl pH 7.5, 150 mM NaCl, 0.5 mM EDTA, 0.01% IGEPAL. For protein– protein interaction assays, the proteins were eluted from the beads with 100 µl buffer (10 mM Tris-HCl pH 8.0, 6 M urea, 2 M thiourea). For kinase activity assays, three more washing steps were carried out, once with 10 mM Tris-HCl, pH 7.5, 300 mM NaCl, 0.5 mM EDTA and twice with 40 mM Tris-HCl pH 7.5, 10 mM MgCl2, 0.1% BSA, 2 mM DTT.

### Kinase activity assay

SIRK1-GFP fusion proteins were affinity purified over anti-GFP beads (see above). A luciferase-based kinase activity assay was performed as described (Wu et al., 2013). The agarose beads with GFP tagged proteins were re-suspended in 30 µl kinase reaction buffer with ATP and the generic kinase substrate myelin basic protein (40 mM Tris-HCl pH 7.5, 10 mM MgCl2, 0.1% BSA, 2 mM DTT, 100 µM ATP, 0.4 µg µL^-1^ myelin basic protein). After incubation, 30 µl ADP-GLO Reagents (Promega) was added. Then, Kinase Detection Reagents were added and incubated for another hour. Luminescence as a measure of ATP conversion from ADP was recorded with a luminometer (TecanM200Pro). Proteins from three independent protein isolations were averaged.

### Binding assays of PEP7-HIS to immobilized SIRK1

The C-terminal HIS-tagged PEP7 was used as bait (Pepmic). 20 μl HisPur™ Ni-NTA Magnetic Beads (Thermo Scientific) were equilibrated 100 mM NaH2PO4/Na2HPO4, 600 mM NaCl, 0.05% Tween™-20, 30 mM imidazole, pH 8.0 and the beads were collected by magnet. The mixture of 100 μl equilibration buffer containing final concentration 1 µM PEP7-HIS was incubated with beads for 1h at 4°C. After incubation, the beads were washed three times with 100 mM NaH2PO4/Na2HPO4, 600 mM NaCl, 0.05% Tween™-20 Detergent, 50 mM imidazole, pH 8.0. SIRK1 full length-GFP protein extract from *N. benthamiana* (50 µg), SIRK1 full length-GFP (50 µg) in microsomal fraction from SIRK1 overexpression line, or purified SIRK1ECD-HA-StrepII (20 µg) were added as prey proteins. Formed complexes were enriched on a magnet and eluted with 40 μl 100 mM NaH2PO4/Na2HPO4, 600 mM NaCl, 250 mM imidazole, pH 8.0. Ten micrograms of the eluate were vacuum-dried and stored for further use.

### Binding assays of SIRK1 to immobilized PEP7-HIS

SIRK1 full length-GFP protein obtained from membrane fraction and SIRK1-ECD-HA-StrepII protein obtained from transient expression in *N. benthamiana* were used as bait to capture PEP7. For binding assay of SIRK1 full length-GFP with PEP7, 100 μg SIRK1-GFP protein extract was prebound to GFP-Traps®-MA beads (Chromotek) for two hours under rotation at 4 °C. The slurry was separated with magnet, the supernatant was discarded and the beads were washed three times with 500 μl of 10 mM Tris- HCl pH 7.5, 150 mM NaCl, 0.5 mM EDTA. Then, prey PEP7 (1 μM) was added to the beads for 1 hour at 4 °C. Beads were washed three more times and the proteins were eluted with 60 μl 6 M urea, 2 M thiourea. For binding assay of SIRK1-ECD to PEP7, 200 μg SIRK1-ECD protein extracted from *N. benthamiana* was prebound to Strep-tag®II beads (Thermo Scientific) for two hours under rotation at 4 °C. The beads were collected by centrifugation and washed three times in 100 mM HEPES pH8.0, 100 mM NaCl, 0.5 mM EDTA, 0.05% Triton X-100 and 2 mM DTT. PEP7 was added and bead slurry was eluted with 100 μl 100 mM HEPES pH 8.0, 100 mM NaCl, 0.5 mM EDTA, 0.05% Triton X-100, 2 mM DTT and 10 mM biotin (Witte et al., 2004). Ten micrograms of the eluates were vacuum-dried and stored for further use.

### Competitive Binding assay

All buffer used were identical to those described for the PEP7-HIS pull-down assay described above. PEP7-HIS (final concentration 1 µM) was firstly immobilized on 20 μl HisPur™ Ni-NTA Magnetic Beads and subsequently 100 μg purified protein SIRK1ECD- HA-StrepII were bound to the immobilized PEP7-HIS. Different concentration of untagged PEP7 (100 μl ddH2O containing 0 μM, 0.2 μM, 0.5 μM, 1 μM, 1.5 μM, 2 μM, 3 μM PEP7) was added to elute the SIRK1-ECD from the immobilized PEP7-HIS. The remaining SIRK1ECD which was not eluted by PEP7 was then eluted with HIS-beads elution buffer (100 mM NaH2PO4/Na2HPO4 pH 8.0, 600 mM NaCl, 250 mM imidazole).

### Microscale thermophoresis

A buffer exchange column was firstly applied to purified SIRK1- ECD protein to remove any unfavorable reagents. The RED-NHS 2^nd^ generation amine reactive dye (MO-L011, NanoTemper Technologies) was used to label the purified SIRK1-ECD (2 µM) with a 5:1 dye:protein ratio for 30 minutes at room temperature in the dark. Column B was employed to remove the excess of dye and produced the labelled protein in PBST buffer (20 mM NaH2PO4/Na2HPO4, 100mM NaCl, pH 7.5, 0.05 % (v/v) Tween 20). Labelled SIRK1-ECD was adjusted to a final concentration of about 20 nM in PBST buffer and then titrated with serial (1:1) dilutions of PEP7 (starting at 0.5 mM), PEP6 or PEP4 (Pepmic). The complex was allowed to establish for 10 min at room temperature before the samples were loaded into the capillaries (MO-K025, NanoTemper Technologies). Binding was detected by a Monolith NT.115 instrument (NanoTemper Technologies) at 24 °C with 80 % excitation power and 40 % MST power, and a dose response curve was generated. QSK1-ECD with a final concentration 20 nM was applied for triple binding affinity estimation. All experiments were repeated at least three times. Raw data was analysed by MO Affinity Analysis software (V2.2.4) and OriginPro software.

### Protoplast swelling assays

Surface sterilized seeds after 48h vernalization were germinated and grown vertically on sucrose starved medium (½MS solid medium with 0.02% sucrose). Approximately 30 seedlings were cut into small pieces and cell walls were digested in 300 mM mannitol, 10 mM MES-KOH pH 5.8, 10 mM CaCl2, 10 mM KCl containing 1% (w/v) cellulase Onozuka R10 (Duchefa) and 1% (w/v) macerozyme R10 (Duchefa). After 3 hours of gentle shaking in the dark at room temperature the protoplasts were prepared *via* a 50 μm nylon mesh filter. Protoplasts were then enriched by centrifugation at 80 g at 4 °C for 10 min, and washed three times with ice-cold wash buffer (300 mM mannitol, 10 mM MES-KOH pH 5.8, 10 mM CaCl2, 10 mM KCl). The protoplasts were finally resuspended in 150 μL wash buffer and stored in the dark on ice for at least 30 min before an experiment was started. In principle, protoplast swelling experiments were performed as described earlier (Sommer et al., 2007; Wu et al., 2013). Approximately 20 µl of the protoplast suspension were pipetted to 200 µL high osmolarity solution (Supplementary Table 4) in a perfusion chamber mounted on the stage of an inverted microscope (DMi8, Leica). Protoplasts were allowed to settle down for 5 minutes. The chamber was perfused with 3 ml of high osmolarity solution to select protoplasts sticking well to the glass bottom of the chamber, then perfused with low osmolarity solution. Solution change in the chamber lasted ca. 15 seconds. A video was recorded for 5 minutes to capture the dynamic change of the protoplasts with the time interval of 3 seconds. Buffers with 30 mM sucrose and/or 1 µM PEP7 treatment were described in Supplementary Table 4. The diameter of the protoplasts was measured directly by using the Leica software LAS X Core, followed by the calculation of volume and surface area at each time point. A regression curve was fitted to the volume change and the maximal slope was obtained from the first derivative of the curve. The maximal water flux density that corresponds to aquaporin activity thus was determined by the maximal slope divided by the protoplast surface area at the corresponding time point. Curve fitting and derivative calculation were performed with OriginPro software.

### FLIM-FRET analysis

The coding sequences of SIRK1 and QSK1 were expressed as C-terminal fluorophore fusions in 2in1 vectors, namely pFRETcg-2in1-CC (Hecker et al., 2015). To obtain GFP-donor only controls, a coding sequence of gentamycin fused to mCherry was used. The binary vectors and p19 as gene silencing suppressor were introduced into *GV3101* and infiltrated into 3-4 week old *N. benthamiana* leaves. After infiltration, the plants were put in darkness for two days. Then 1 µM PEP7 or water was added to the system by infiltration into the leaves right before the measurement. In some experiments, the proteinase inhibitor leupeptin was infiltrated directly with the constructs at a concentration of 10 µM. The measurements were performed for a maximum time of 10 min based on a modified protocol (Glöckner et al., 2020) with a SP8 confocal laser scanning microscope (Leica Microsystems) equipped with Leica Microsystems Application Suite software and a FastFLIM upgrade from PicoQuant consisting of Sepia Multichannel Picosecond Diode Laser, PicoQuant Timeharp 260, TCSPC Module and Picosecond Event Timer (Picoquant). Imaging was done by using a ×63/1.20 water-immersion objective and focusing on the plasma membrane of the abaxial epidermal cells. The presence of the fluorophores was detected by excitation with 488 nm or 561 nm, and 500-500 nm or 600-650 detection range for GFP or mCherry, respectively. Colocalization was demonstrated by reading out signal intensities over the plasma membrane. GFP fluorescence lifetime in nanoseconds of either donor-only expressing cells or cells expressing the indicated combinations was measured with a pulsed laser as an excitation light source of 470 nm and a repetition rate of 40 MHz. The acquisition was performed until 500 photons in the brightest pixel were reached at a resolution of 256 x 256 pixel. For data processing a region of interest at the plasma membrane was defined in the SymPhoTime software and bi-exponential curve fitting as well as correction for the instrument response function was applied. A total range of 23 ns was evaluated. Statistical analysis was carried out with JMP 14 and OriginPro software.

### Trypsin digestion and phosphopeptide enrichment

Protein pellets were solubilized in 10mM Tris-HCl pH8.0, 6 M urea, 2M thiourea and by subjecting the protein solution to ultrasonic bath for 10 minutes and in solution trypsin-digested was carried out as described (Wu et al., 2017). Digested peptides were resuspended in 1 M glycolic acid, 80% v/v Acetonitrile and 6% v/v TFA 80%. Phosphopeptides were enriched by TiO2-beads (Titansphere, 5 µm, GL Sciences) as described (Wu et al., 2017).

*LC-MS/MS analysis of peptides and phosphopeptides* – Peptides mixtures were analyzed by nanoflow Easy-nLC (Thermo Scientific) and Orbitrap hybrid mass spectrometer (Q-exactive HF, Thermo Scientific). Peptides were eluted from a 75 µm x 25 cm analytical C18 column (PepMan, Thermo Scientific) on a linear gradient running from 4% to 64% acetonitrile over 135 min. Proteins were identified based on the information-dependent acquisition of fragmentation spectra of multiple charged peptides. Up to twelve data-dependent MS/MS spectra were acquired for each full-scan spectrum acquired at 60,000 full-width half-maximum (FWHM) resolution.

### Peptide and protein identification

Protein identification and ion intensity quantitation was carried out by MaxQuant version 1.5.3.8 (Cox and Mann, 2008). Spectra were matched against the *Arabidopsis* proteome (TAIR10, 35386 entries) using Andromeda (Cox et al., 2011). Thereby, carbamidomethylation of cysteine was set as a fixed modification; oxidation of methionine as well as phosphorylation of serine, threonine and tyrosine was set as variable modifications. Mass tolerance for the database search was set to 20 ppm on full scans and 0.5 Da for fragment ions. Multiplicity was set to 1. For label-free quantitation, retention time matching between runs was chosen within a time window of two minutes. Peptide false discovery rate (FDR) and protein FDR were set to 0.01, while site FDR was set to 0.05. Hits to contaminants (e.g. keratins) and reverse hits identified by MaxQuant were excluded from further analysis. The identified phosphopeptides including their spectra were submitted to the phosphorylation site database PhosPhAt (Durek et al., 2010; Heazlewood et al., 2008). The mass spectrometry proteomics data have been deposited to the ProteomeXchange Consortium *via* the PRIDE (Deutsch et al., 2017) partner repository with the dataset identifiers PXD029498 (affinity purification combined with mass spectrometry experiments), PXD029050 (binding assays), PXD029509 (phosphoproteomics).

### Label-free peptide and protein quantitation

We followed a label-free quantitation approach bases on the LFQ values as quantitative information obtained from MaxQuant (Cox et al., 2014). For protein identification and quantitation, protein groups information (protein groups.txt) was used. For quantitation of phosphopeptides the phosphosite data (Phospho(STY)Sites.txt) was used as they were written by MaxQuant. Data analysis and multivariate statistics was performed by Perseus (Tyanova et al., 2016).

### Statistical analyses and data visualization

Functional classification of proteins was done based on MapMan (Thimm et al., 2004). Subcellular location information was derived from SUBA (Tanz et al., 2013). Protein function was manually updated with TAIR (Poole, 2007). Other statistical analyses were carried out with SigmaPlot (version 11.0) and Excel (Microsoft, 2013).

## Acknowledgements

We thank Julian Ams and Sonja Pressmar for their work on apoplasmic proteomes. We thank Prof. Gerhard Obermeyer and Prof. Heidi-Pertl Obermeyer for advice of establishing protoplast swelling assay system. Research in our laboratories was supported by the German Research Foundation (DFG) with grants to WS (SCHU1533/10-1; SCHU1533/11-1) and to KH (CRC 1101-D02, HA 2146/22-2, HA 2146/23-1) and a grant for scientific equipment (INST 37/819-1 FUGG)

## Supplementary Materials

**Figure S1:**
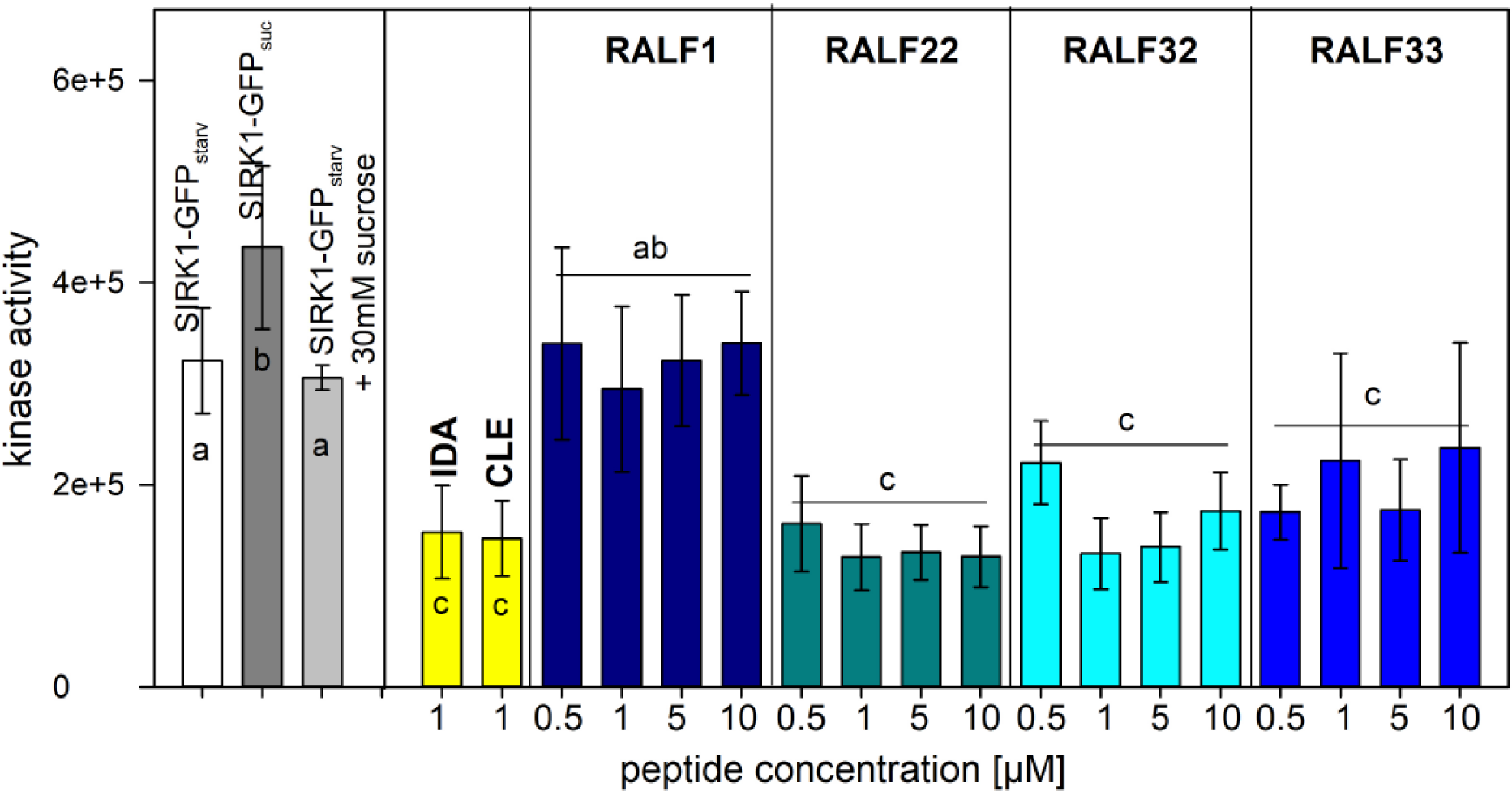
Kinase assay activity assay for SIRK1-GFP enriched from sucrose-starved roots by exposure to different unrelated signaling peptides. IDA and CLE were used as negative controls. RALF peptides were supplied at different concentrations. Averages of three independent assays are shown with standard deviation. Asterisks indicate significant differences to SIRK1-GFP under sucrose starvation (white bar).

**Figure S2:**
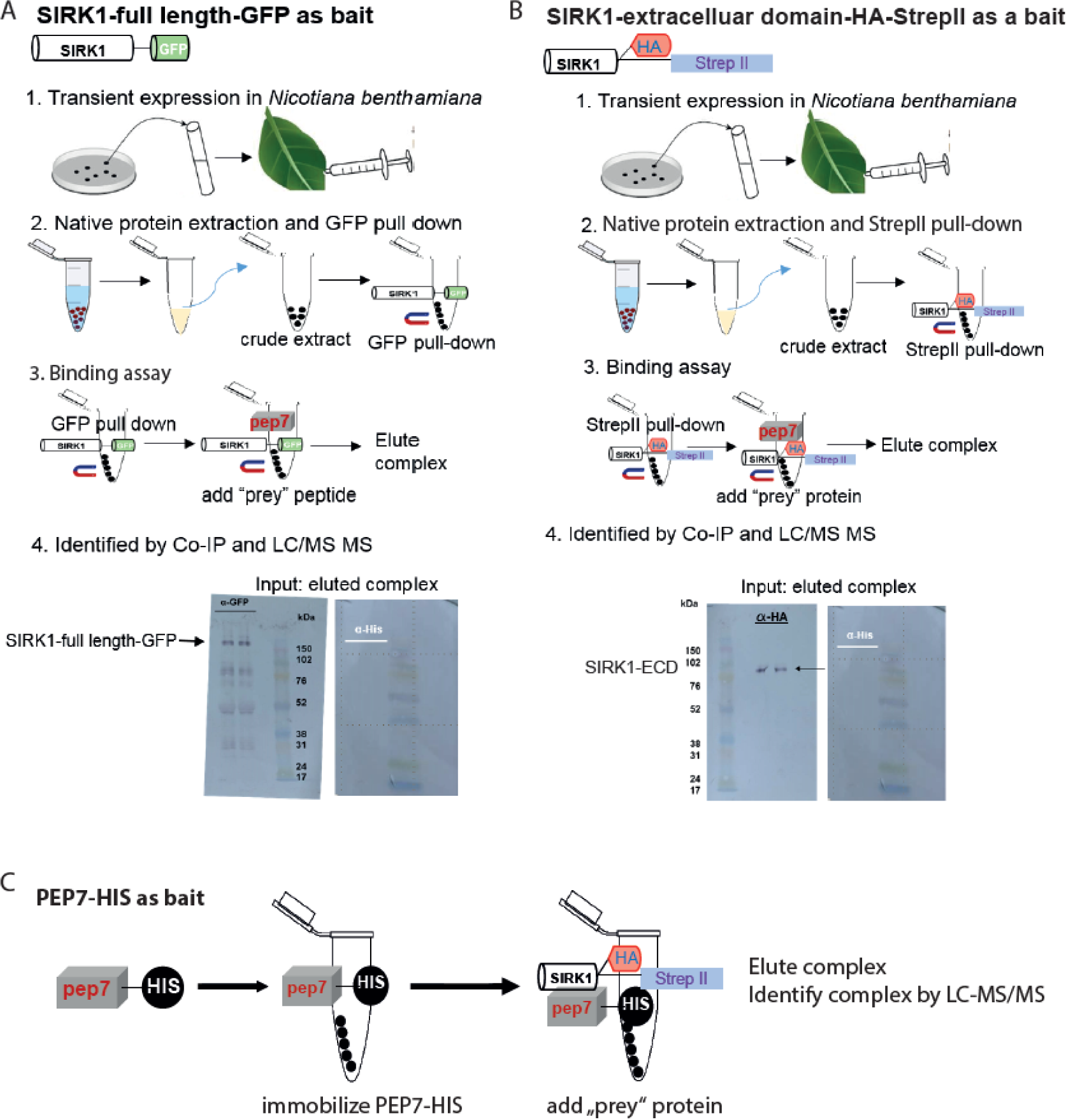
Co-immunoprecipitation workflows to test the interaction of SIRK1 and PEP7. (**A**) Immobilization of SIRK1-GFP to interact with free PEP7. (**B**) Immobilized SIRK1 extracellular domain to interact with free PEP7. (**C**) Immobilized PEP7-HIS6 to interact with SIRK1-GFP, SIRK1 extracellular domain, or native SIRK1 purified from roots.

**Figure S3:**
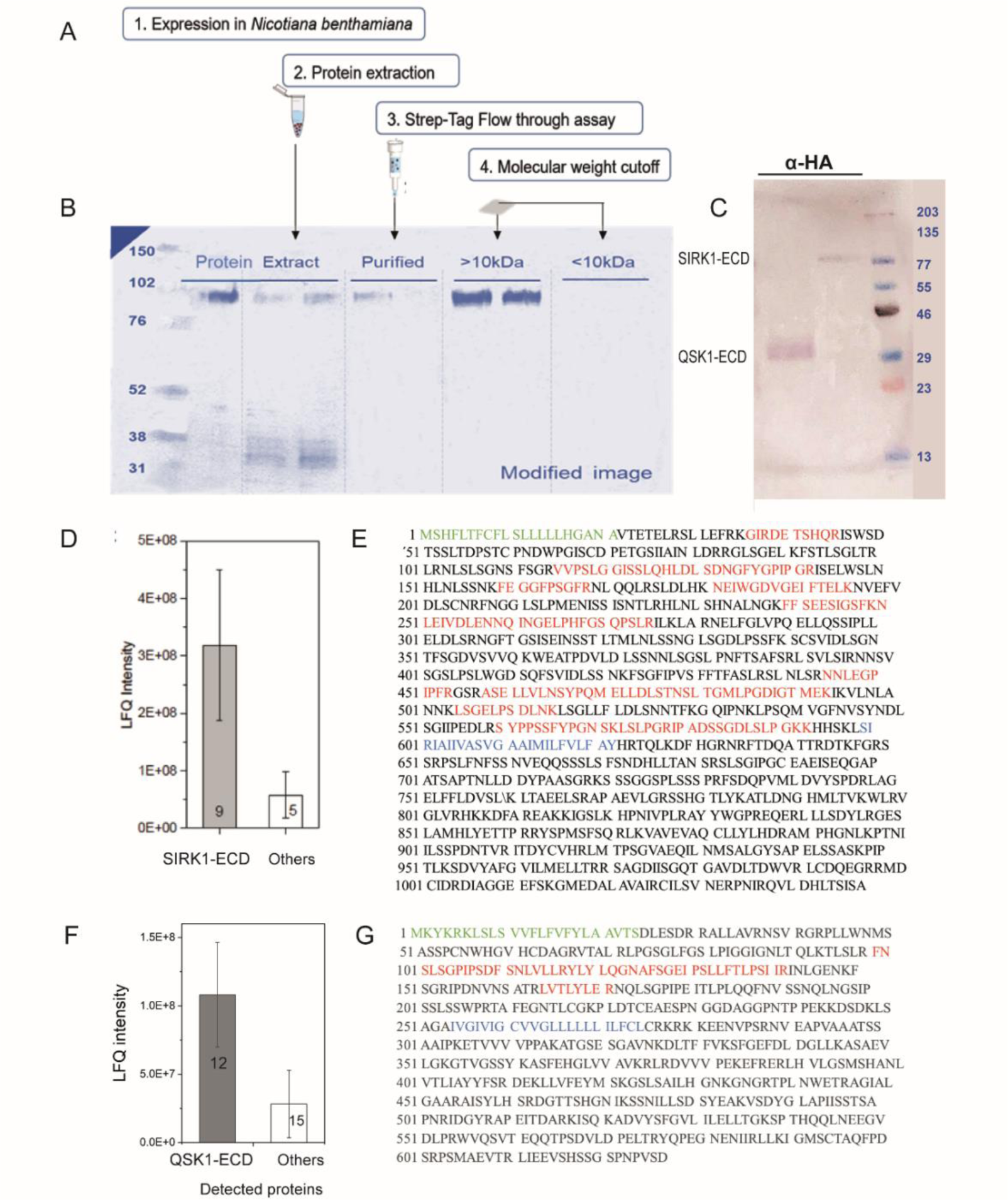
Purification and identification of SIRK1-ECD and QSK1-ECD (**A**) Purification steps of SIRK1-ECD. (**B**) SDS-PAGE analysis of fractions after purifications. (**C**) Western blot of StrepII purified fraction by HA antibody. (**D**) LFQ intensity comparison of tryptic digest peptides from final purified SIRK1-ECD. Nine of the fourteen peptides identified belong to SIRK1, and their summed LFQ intensity was approximately 6-fold higher than the summed LFQ intensity of all the other identified peptides. (**E**) Tryptic digest peptides mass fingerprint analysis of purified SIRK1-ECD. (**F**) LFQ intensity comparison of tryptic digest peptides from final purified QSK1-ECD. (**G**) Tryptic digest peptides mass fingerprint analysis of purified QSK1-ECD. The sequence of full length SIRK1 and QSK1 is shown with signal peptide in green and the transmembrane domain in blue. All identified peptides from the sample were shown in red. All identified SIRK1 peptides cover only the extracellular domain of SIRK1, so does QSK1. Taken together, the purified SIRK1-ECD was considered to meet quality criteria for fluorescent labelling and MST assays.

**Figure S4:**
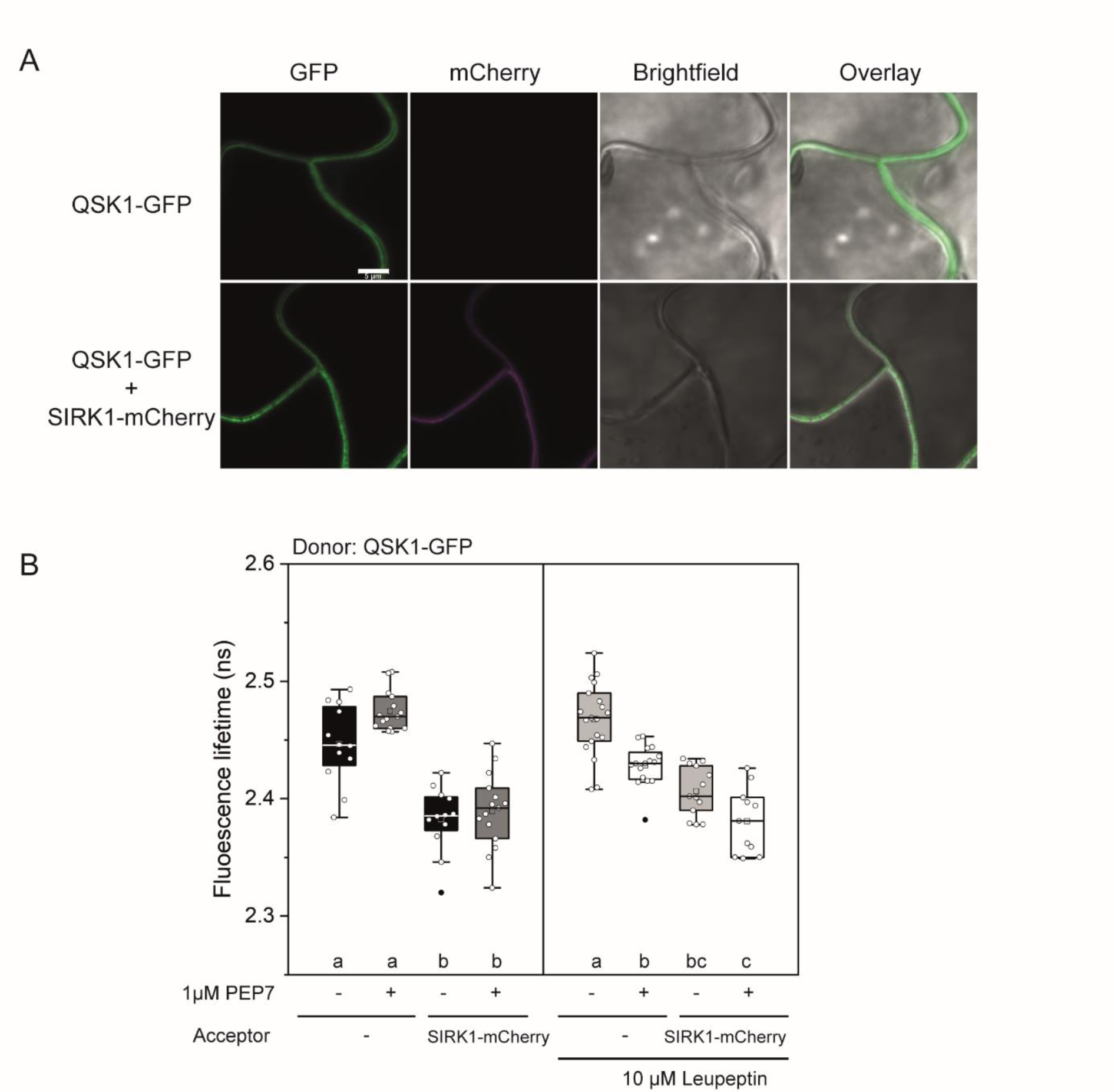
FRET-FLIM of SIRK1 and QSK1 in presence and absence of PEP7. **(A)** Representative confocal images show co-localization of SIRK1 and QSK1 with or without infiltration of 1 µM PEP7. Constructs were transiently transformed in *N. benthamiana* leaf cells. Scale bars indicate 5 μm. **(B)** FRET-FLIM reveals spatial proximity of SIRK1 and QSK1 in presence and absence of proteinase inhibitor leupeptin. For each of measurements, n ≥ 7. Statistical differences in FLT were analysed using a two-way ANNOVA followed by Dunn-Sidak multiple comparison test. Small letters indicate significantly different means at p<0.05. The centre line indicates the median, the bounds of the box show the 25^th^ and the 75^th^ percentiles, and the whiskers indicate 1.5 × IQR.

**Figure S5:**
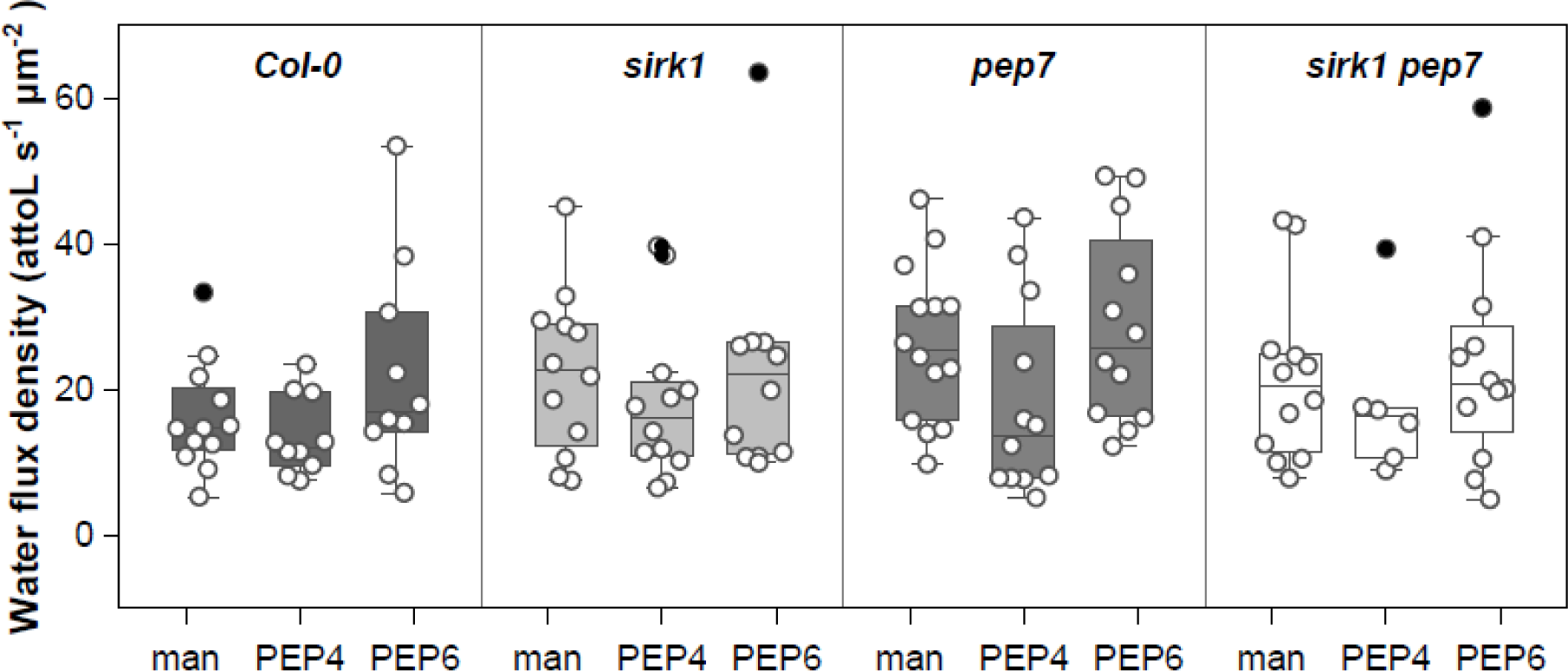
Water influx rates of protoplasts with and without supply of PEP4 and PEP6. Boxplots show the median and upper/lower 25^th^ percentile. White dots represent individual measurements. Man, mannitol.

**Figure S6.**
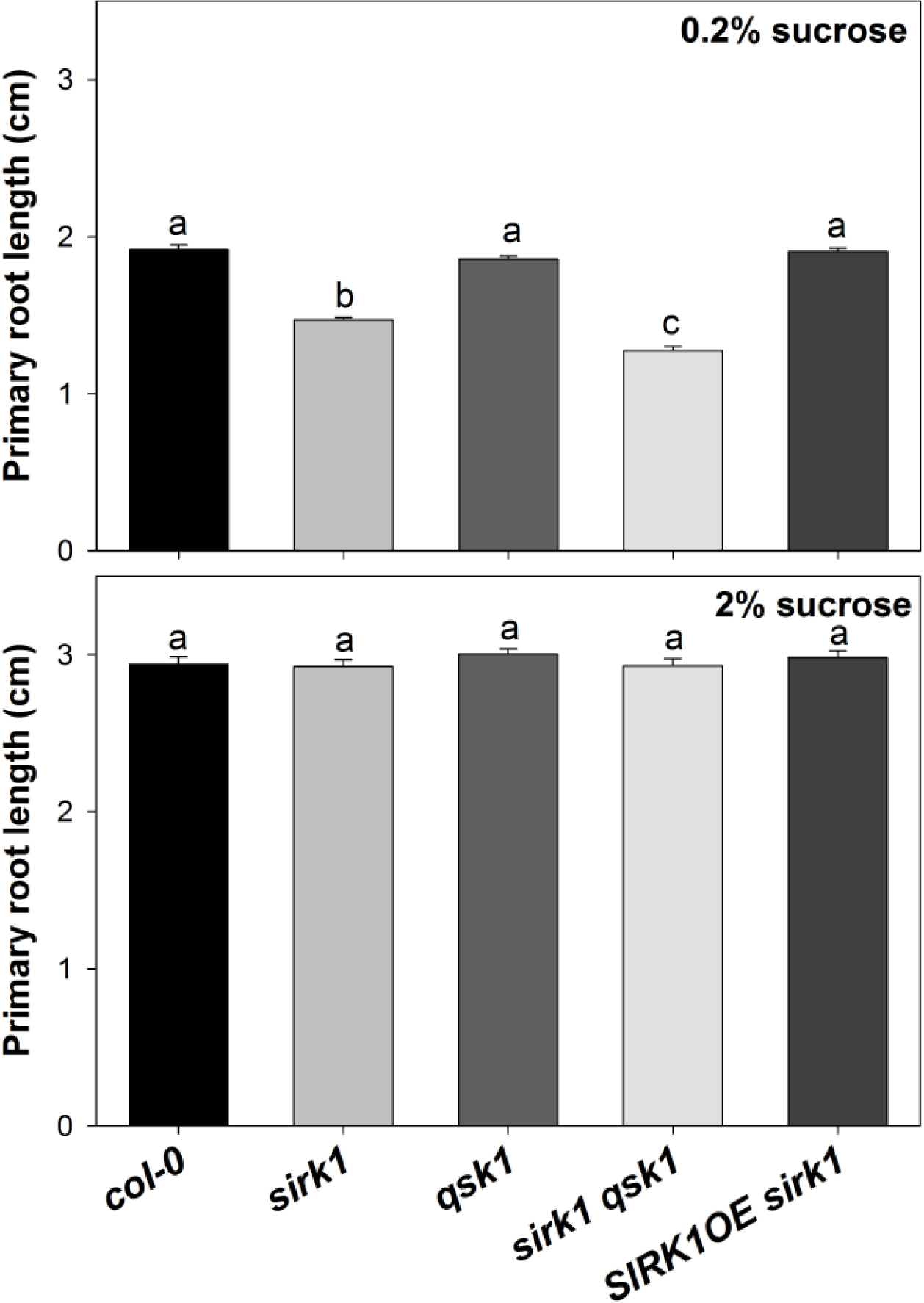
Root growth of wild type (Col-0) *sirk1*, *qsk1*, *sirk1 qsk1*, as well as *SIRK1-OE sirk1* at (**A**) low and (**B**) high external sucrose. Bar plots indicate measured primary root lengths of at least 30 seedlings with standard deviations. Small letters indicate significant differences (p<0.05, t-test) within one condition compared to wild type.

**Figure S7:**
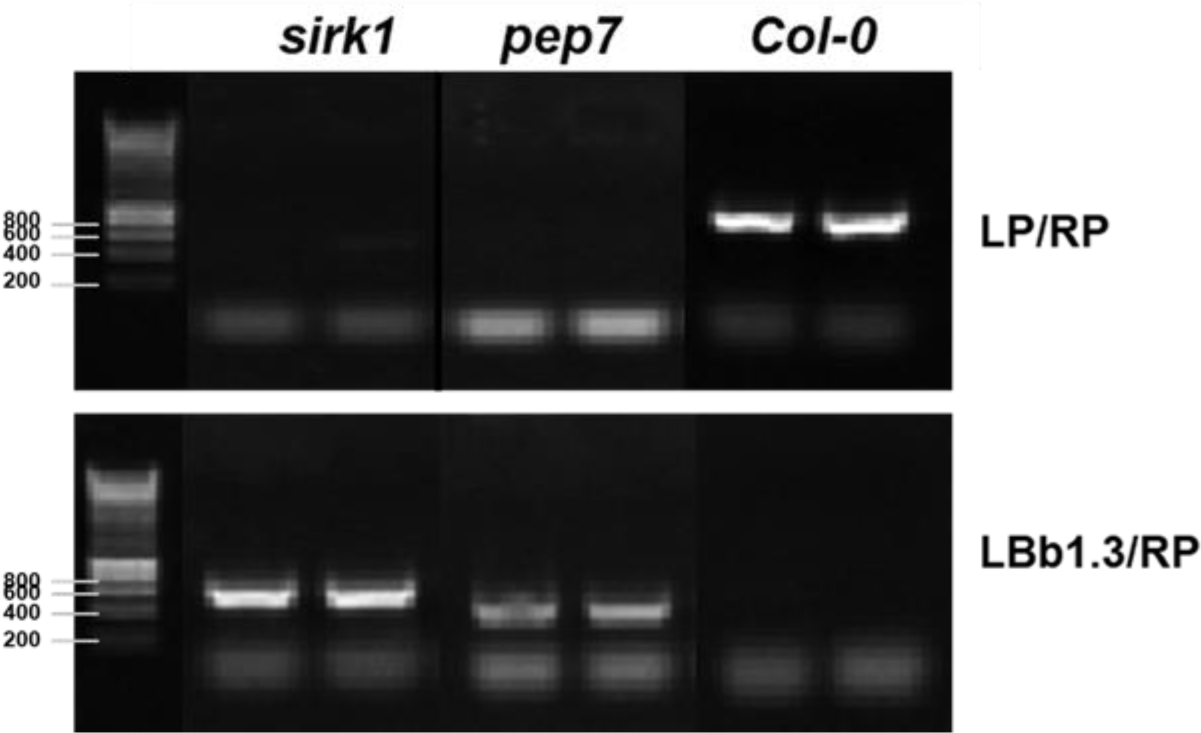
Identification of double mutant of *sirk1pep7* as a result of crossing single mutants *sirk1* (SALK_125543) and *pep7* (SALK_025824).

## Supplementary Tables

Supplementary Table 1: Overview of the proteins identified in tryptic and non-tryptic samples in fractions F1, F2 and F3.

Supplementary Table 2: 1076 common Interaction partners of SIRK1 under different treatments.

Supplementary Table 3: List of identified phosphopeptides from which the heatmap in Figure 4 was generated.

**Supplementary Table 4:**
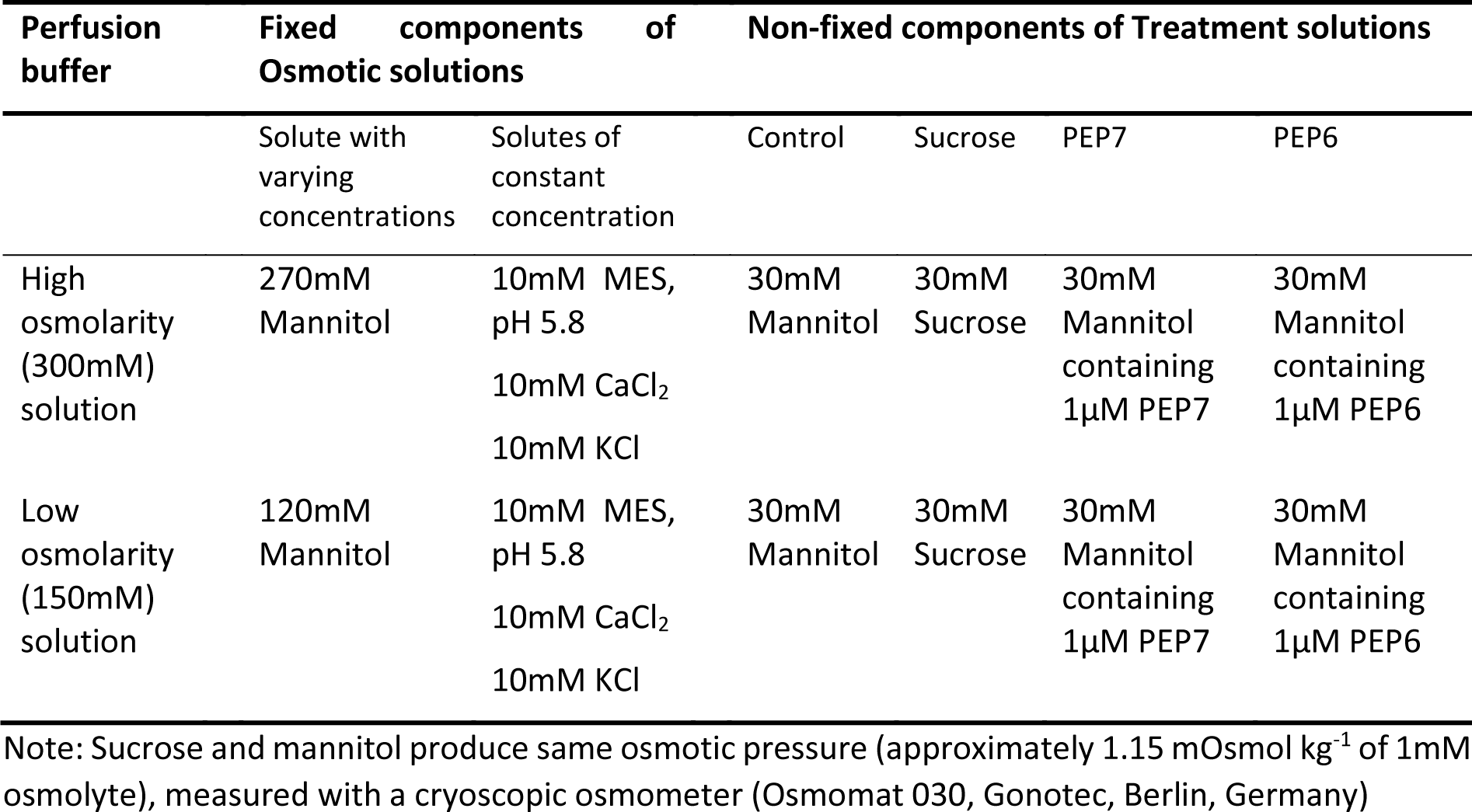
Overview of the buffers used in the protoplast swelling assays.

